# Identification of SARS-CoV-2 induced pathways reveal drug repurposing strategies

**DOI:** 10.1101/2020.08.24.265496

**Authors:** Namshik Han, Woochang Hwang, Konstantinos Tzelepis, Patrick Schmerer, Eliza Yankova, Méabh MacMahon, Winnie Lei, Nicholas M Katritsis, Anika Liu, Alison Schuldt, Rebecca Harris, Kathryn Chapman, Frank McCaughan, Friedemann Weber, Tony Kouzarides

**Affiliations:** Milner Therapeutics Institute, University of Cambridge, Cambridge, UK; Institute for Virology, FB10-Veterinary Medicine, Justus-Liebig University, Gießen 35392, Germany; Centre for Therapeutics Discovery, LifeArc, Stevenage, UK; Department of Chemical Engineering and Biotechnology, University of Cambridge, Cambridge, UK; Department of Chemistry, University of Cambridge, Cambridge, UK; Department of Medicine, University of Cambridge, Cambridge, UK; The Gurdon Institute and Department of Pathology, University of Cambridge, Cambridge, UK

## Abstract

The global outbreak of SARS-CoV-2 necessitates the rapid development of new therapies against COVID-19 infection. Here, we present the identification of 200 approved drugs, appropriate for repurposing against COVID-19. We constructed a SARS-CoV-2-induced protein (SIP) network, based on disease signatures defined by COVID-19 multi-omic datasets(Bojkova et al., 2020; Gordon et al., 2020), and cross-examined these pathways against approved drugs. This analysis identified 200 drugs predicted to target SARS-CoV-2-induced pathways, 40 of which are already in COVID-19 clinical trials(Clinicaltrials.gov, 2020) testifying to the validity of the approach. Using artificial neural network analysis we classified these 200 drugs into 9 distinct pathways, within two overarching mechanisms of action (MoAs): viral replication (130) and immune response (70). A subset of drugs implicated in viral replication were tested in cellular assays and two (proguanil and sulfasalazine) were shown to inhibit replication. This unbiased and validated analysis opens new avenues for the rapid repurposing of approved drugs into clinical trials.

## INTRODUCTION

To date, the majority of small molecule and antibody approaches for treating SARS-CoV-2 related pathology are rightly rooted in repurposing and are focused on several key virus or host targets, or on pathways as points for therapeutic intervention and treatment. This has been underpinned by the unprecedented pace of scientific research to uncover the molecular bases of virus structure, and the mechanisms by which it gains access to cells before replication and release of new virus particles. The emergence of global proteomics datasets is now propelling our understanding of these mechanisms through which the virus interacts with host cell proteins, determining the Directly Interacting Proteins (DIP)(Gordon et al., 2020) and Differentially Expressed Proteins (DEP)(Bojkova et al., 2020). Such interactome outputs, and related efforts in transcriptomics(Blanco-Melo et al., 2020), have begun to provide detailed information on possible individual targets and pathways against which currently available drugs can be tested for potential COVID-19 repurposing. Systematic analyses of these datasets will direct further research to likely points of successful therapeutic intervention. In this study, we have applied the power of bespoke computational biology and machine learning approaches to dissect these datasets and construct an agnostic network for SARS-CoV-2-induced pathways, uncovering novel targets and potential repurposing strategies.

## RESULTS

### Construction of a SARS-CoV-2-induced protein (SIP) network

To determine the disease mechanisms underlying a SARS-CoV-2 infection, we undertook a comprehensive analysis of the protein pathways implicated in COVID-19, using computational biology workflows for data integration and network construction. To this end, we hypothesized that DIP are the ‘cause’ and DEP are the ‘consequence’ of SARS-CoV-2 infection. We then constructed a fully connected SIP network between DIP and DEP to understand how the chains of the cause and consequences are connected (Figure 1A). In our SIP network, there are three layers: the DIP, the DEP and the hidden layer between the two. To make a fully connected network between the DIP and DEP, we identified all possible shortest paths between these. Our analysis of the DEP data identified differentially expressed proteins at 6-hours and 24-hours after infection; hence we constructed the SIP network for these two time points. There are 13,308 proteins and 344,543 interactions in the 6-hour network and 14,827 proteins and 528,969 interactions in the 24-hour network (Figure S1 shows the entire SIP network at the two timepoints, Figure 1B shows a sub-network of SARS-CoV-2 Orf8 at 24-hour). Among the DIP-to-DEP paths in both networks, ~99% of paths are via more than one protein (Figure 1C) which suggests that the ‘hidden layer’ we have constructed in our network is central to understanding the pathways that connect DIPs and DEPs.

**Figure 1.**
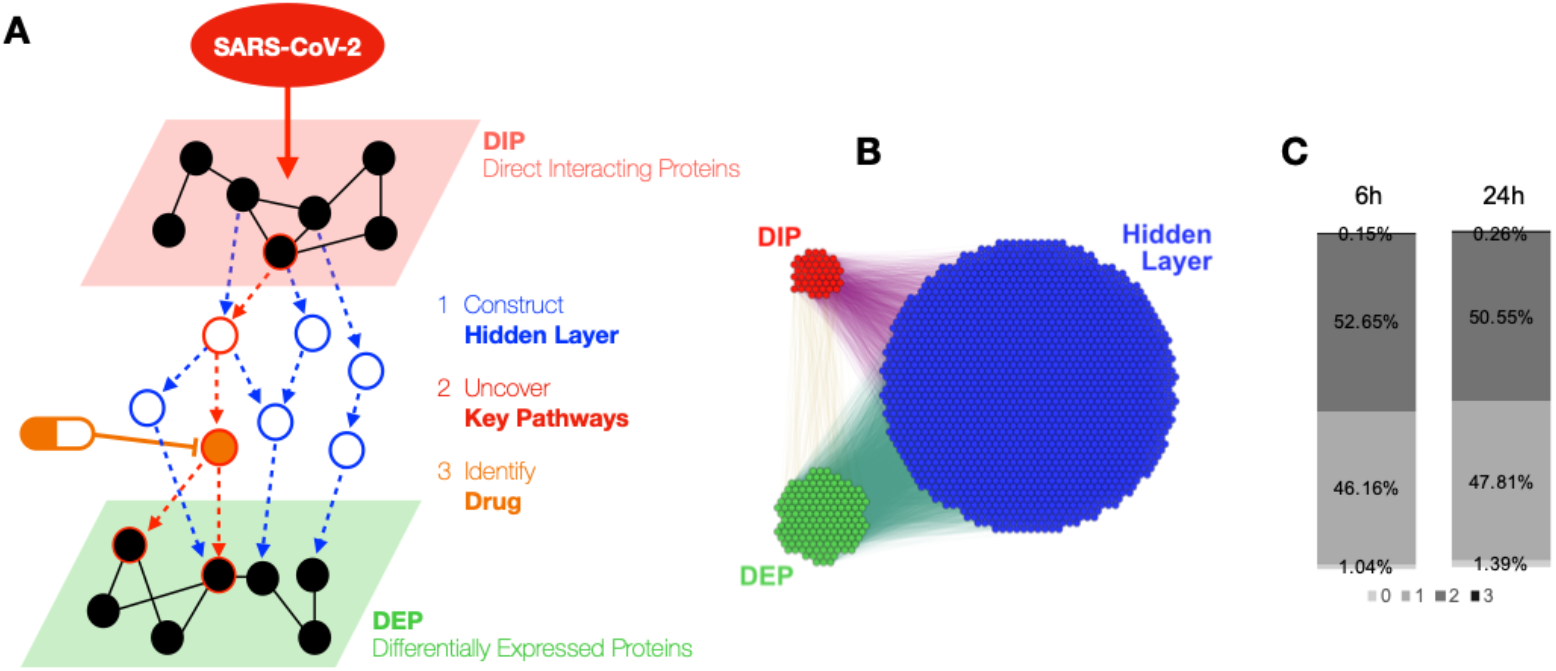
Construction of a SARS-CoV-2-induced protein (SIP) network identifies key pathways. (A) The schematic depicts our strategy of constructing a SARS-CoV-2-induced protein (SIP) hidden network through data integration and network construction of directly-interacting proteins (DIPs) and differentially expressed proteins (DEPs), followed by identification of drugs that target key pathways in this network (B) The SARS-CoV-2 Orf8 sub-network shows the extent of the hidden layer that is revealed through the network analysis (C) Percentage of shortest paths between the DIP and DEP that are via 0-3 proteins at 6h versus 24h.

### The SIP network can be interrogated to reveal key proteins and disease pathways

To identify important proteins and key pathways in the SIP network, we applied multiple network algorithms and a permutation test to identify statistically significant proteins (see Methods). This revealed 320 proteins at 6h and 394 proteins at 24h, of which 238 (50% of 476 proteins) proteins were in common (Figure 2A). More than half of the proteins identified as significant at both time points were in the hidden layer: 170 (53%) and 202 (51%), respectively (Figures S2A and S2B). We then asked whether these proteins were also biologically relevant to the disease symptoms caused by COVID-19. A disease enrichment analysis on the proteins showed that the top 10 enriched diseases for these proteins at both timepoints are diseases that are potentially relevant for COVID-19 pathogenesis, including lung disease(Guan et al., 2020; Gupta et al., 2020), hypertension(Guan et al., 2020; Gupta et al., 2020) and hyperglycaemia(Gupta et al., 2020) (Table S1). To uncover potential biological functions of the important proteins at 6h, 24h and both timepoints, we took a closer look at key pathways in which they are enriched. For proteins at 6h and proteins that are common to both timepoints, the pathways were related to the immune system and virus replication (Figures S3A and S3B). In contrast, the pathways that were relevant for the proteins at 24h were primarily related to virus replication (Figure S3C). In this way, we established a COVID-19 SIP network that allows investigation of disease pathways that are pertinent to SARS-CoV-2 infection.

**Figure 2.**
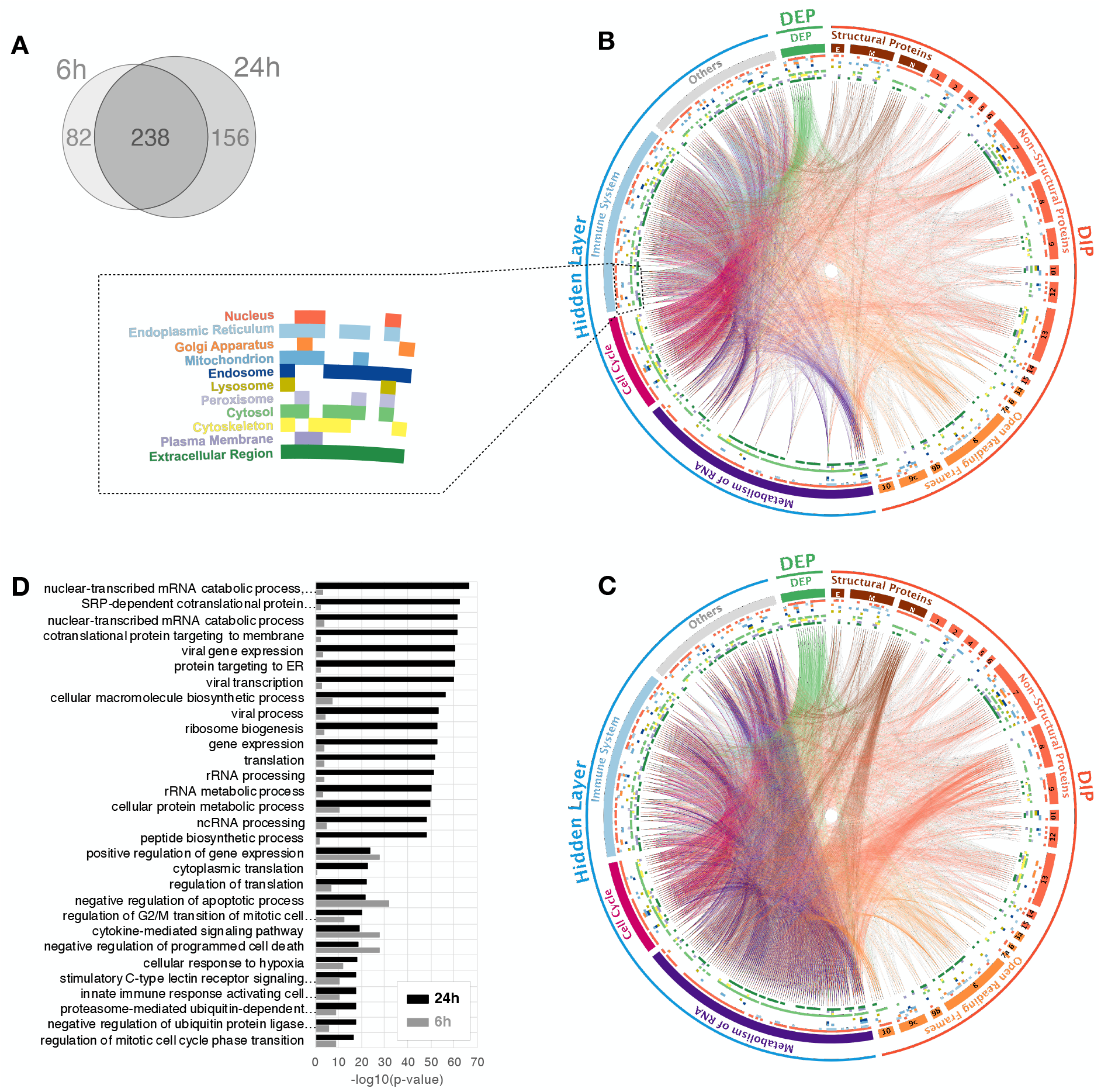
Sub-network analysis shows an enrichment of viral replication pathways. (A) Venn diagram of key proteins in 6-hour and 24-hour SIP networks (B) A circos plot depicting interactions between DIPs and DEPs revealed through the SIP network at the 6h after infection. DIPs were sub-divided into the genomic organization of SARS-CoV-2. Proteins in the hidden layer were also sub-divided into major pathways. Inner coloured circles demonstrate the subcellular localization of the proteins, and details are shown in the dotted box. The coloured lines show protein-protein interaction. (C) and 24h after infection (D) GO term enrichment for the pathways in the SIP network at 6h and 24h.

### Sub-network analysis demonstrates an enrichment of pathways related to viral replication

To understand the disease mechanism of COVID-19, we investigated which biological processes SIP sub-networks are implicated in, for 29 SARS-CoV-2 proteins (4 structural proteins, 16 non-structural proteins and 9 accessory factors of the virus genome). We analysed several parameters: (1) the subcellular localization of the proteins; (2) the differences between the 6h and 24h timepoints (Figures 2B and 2C); and (3) the biological processes that the proteins act in. We found significantly stronger relevance of RNA metabolism at 24h. We observe that the viral proteins N (Nucleocapsid), Nsp 8 (Non-structural protein 8), Orf 8 (Open reading frame 8) and Orf 10 of SARS-CoV-2 interact with ribosomal proteins in the hidden layer of our SIP network, indicating that they may have a possible influence on RNA metabolism. The N and Nsp 8 proteins are known to drive viral replication(Gordon et al., 2020). More interestingly, Orf8 and 10 are the only two proteins of SARS-CoV-2 that are distinct from other coronaviruses(Tang et al., 2020). We observed that Orf8 was enriched in the endoplasmic reticulum (ER) (Figure 2C), which may be significant as the ER is the intracellular niche for viral replication and assembly(Romero-Brey and Bartenschlager, 2016). There were 54 key proteins in the hidden layer that did not have strong enrichment in known biological pathways (‘Other’) and that actively interacted with ‘virus replication’ proteins at 24h. Further study on the unknown proteins found individual links to RNA binding (ATP5A1, MRTO4 and NHP2L1), host-virus interaction (ACE2, CXCR4, DERL1, GNB2L1, HSPD1, KDR, KRT18, SIRT1 and TMPRSS2), histones (H2AFZ, HIST2H3PS2 and WDTC1), viral mRNA translation (MRPS7) and ER-associated responses (ATF4, CFTR, DERL1 and INS). We next confirmed statistically that virus-related pathways are enriched in the top enriched GO-terms as well as RNA- and ER-related processes (Figure 2D). The differences between two time points were also confirmed. In summary, our pattern analysis in the SIP sub-networks revealed biological pathway changes during the course of infection, with prominent increases in proteins involved in virus replication by 24h.

### An *in silico* drug simulation on the key pathway of SIP network identifies drug candidates

To identify drugs targeting the key pathways, we conducted a network-based *in-silico* drug efficacy simulation(Guney et al., 2016) on the key pathways of SIP network at 6h and 24h after infection. We collected 1,917 approved drugs from publicly available databases (ChEMBL(Mendez et al., 2019) and DrugBank(Wishart et al., 2018)). This virtual screening identified 200 drugs (Table S2) that are predicted to target the key pathways of SIP network, of which 99 (49.5%) were specific to the 6h timepoint, 14 (7%) were specific to the 24h timepoint and 87 (43.5%) were common to both timepoints. We then checked the Anatomical Therapeutic Chemical (ATC) code (available for 180 drugs only) to determine the therapeutic areas for which specific drugs have been developed. The top clinical areas against which these approved drugs are used for were cancer, sex hormone signalling, diabetes, immune system, bacterial disease and inflammatory/rheumatic disease (Figure S4). Interestingly, 35% of the 200 drugs have been tested in phase 2 or 3 clinical trials for infectious diseases, and half of these were HIV trials; further drugs have been tested in trials for inflammatory (16%) and respiratory (10%) disease.

Among the 200 identified drugs, 40 (20%) are now in COVID-19 clinical trials(Clinicaltrials.gov, 2020) (Table S3). To determine the significance of this finding, we asked what the likelihood would be of this number of drugs being identified as hits by chance and found that, by comparison, only 249 of 1,917 drugs were in the COVID-19 clinical trials(Clinicaltrials.gov, 2020). A hypergeometric test returned a p-value of 3.59e-03, demonstrating the reliability and validity of our computational approaches. 30 further drugs that we identified have also been reported as being potential candidates against COVID-19(Courtney J. Mycroft-West, Dunhao Su, Yong Li, Scott E. Guimond, Timothy R. Rudd and Elli, Gavin Miller, Quentin M. Nunes, Patricia Procter, Antonella Bisio, Nicholas R. Forsyth, Jeremy E. Turnbull, Marco Guerrini, David G. Fernig, Edwin A. Yates, 2020; Criado et al., 2020; Kuleshov et al., 2020; kumar, arun; C.S, Sharanya; J, Abhithaj; C, 2020; MirIbrahim Sajid, Javeria Tariq, SheharBano Awais, ZehraNaseem, SamiraShabbir Balouch, 2020; Shin et al., 2020). Thus, drug efficacy simulation has revealed 70 drugs in total that are either in COVID-19 clinical trials or being considered as potential drug candidates in pre-clinical studies, supporting the strength of our approach. In addition, we have discovered 130 additional drugs that could provide novel opportunities for repurposing as COVID-19 therapeutics. The full list of 200 approved drugs along with their safety profile and MoA are shown in Table S2.

### Artificial neural network analysis uncovers drug mechanisms of action

To investigate the mechanism of action (MoA) for the 200 drugs in the context of COVID-19, we used Self-Organizing Map (SOM), a type of artificial neural network, to analyse the relationship between the 200 drugs and the 148 key pathways (termed drug-pathway association). After the unsupervised training of SOM, the distance between the adjacent neurons (pathways) was calculated and presented in different coloured hexagons, which illustrates the probability density distribution of data vectors (drug-pathway association score) (Vesanto and Alhoniemi, 2000) (Figures 3A and S5). Based on the distance, we applied the Davies-Bouldin (DB) index to separate the key pathways into 9 clusters (Figure 3B). These clusters of pathways and drugs identified two MoA categories: (1) virus replication (VR) and (2) immune response (IR) (Figure 3C). The SOM also mapped 200 drugs into each neuron (the number of drugs per neuron is shown in Figure 3D and drug names are shown in Figure 3E). Notably, 30 out of the 40 drugs that are in COVID-19 clinical trials(Clinicaltrials.gov, 2020) were in the VR MoA category while only 10 drugs were in the IR (Figure 3D). Finally, we identified mechanistic roles and connections for the 200 drugs and their target proteins, and mapping the drugs into 9 pathway clusters (Figure 3E). A more extensive analysis of information about each drug is given in Table S2.

**Figure 3.**
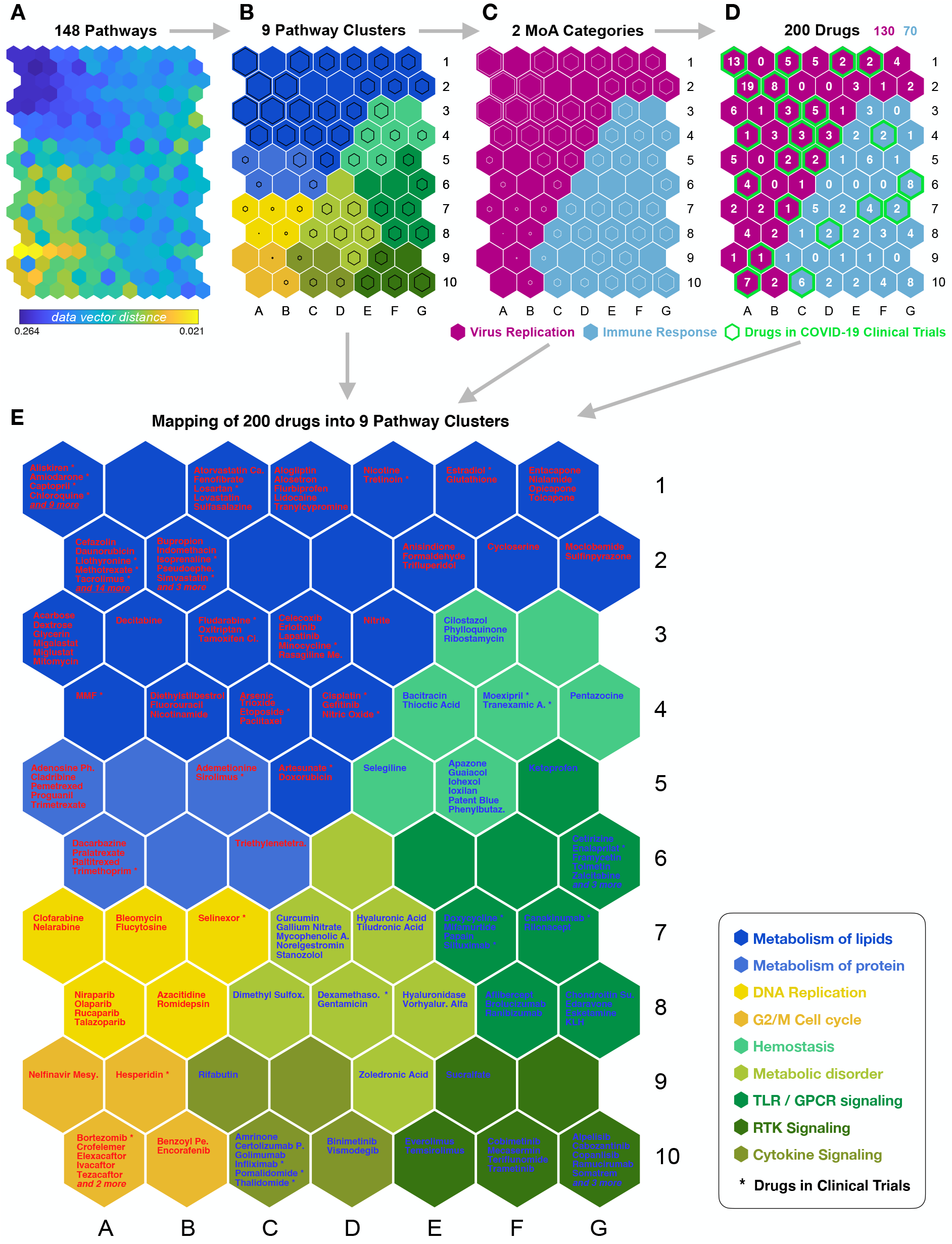
Machine learning predicts mechanisms of actions for drug repurposing candidates. (A) A unified distance matrix (U-matrix) is shown of the trained unsupervised SOM used to analyse the relationship between the 200 drugs and the 148 key pathways. This contains the distance (similarity) between the neighbouring SOM neurons (pathways) and shows data density (drug-pathway association scores) in input space. Each subunit is coloured according to distance between corresponding data vectors of neighbour neurons, with low distances areas (dark blue) indicating high data density (clusters). (B) Cluster solution chosen based on U-matrix and Davies-Bouldin (DB) index to separate the key pathways into 9 clusters. Clusters of each SOM neuron are distinguishable by colour. The size of the black hexagon in each neuron indicates distance. Larger hexagons have a low distance to neighbouring neurons, hence forming a stronger cluster with neighbours. (C) Two MoA categories identified based on the pathway clustering and the drug mapping. (D) Mapping of the 200 identified drugs to each neuron (pathway) based on matching rates and inspection of examples from each cluster. (E) A SOM component map shows mapping results of the 200 drugs into 9 pathway clusters. The names of 9 clusters are shown in the figure, and the drugs with asterisk are already in COVID-19 clinical trials.

We next sought to identify the precise proteins, within the SIP network, targeted by each of the 200 drugs. We found that of the 1,573 proteins targeted by the 200 drugs, most (66%) are targeted by a single drug (Figure S6A). However, there are 30 proteins (0.19%) that are targeted by 8 or more drugs (Figure S6A). To establish whether there is a pathway relationship between these 30 proteins, we interrogated their molecular function. Figure S6B shows that the most enriched categories of function for these proteins were heme, microsome, oxidoreductase and monooxygenase, all of which are related to nicotinamide adenine dinucleotide phosphate (NADP) and nitric oxide (NO) synthesis. As NO is important for viral synthesis (and because NADP affects NO production), this could provide a potential mechanism by which these drugs might alter viral infection(Kwiecien et al., 2014; Lind et al., 2017; Wang et al., 2006). Based on these findings we decided to validate in cellular assays, five drugs (Ademetionine, Alogliptin, Flucytosine, Proguanil and Sulfasalazine) with good safety profiles which are functioning within this pathway.

### Two drugs that target NO production reduce SARS-CoV-2 replication

To assess whether these five drugs are able to reduce SARS-CoV-2 infection, we performed an initial screening using the Vero E6 cell line, where we observed that 2 of the 5 drugs, Proguanil and Sulfasalazine, showed significant antiviral effects without any noticeable cellular toxicity at the indicated doses (Figures 4A and S7A). We then focused on these two drugs, expanding our validation using 2 different cellular models (Vero E6 and Calu-3). Treatment of Vero E6 and Calu-3 cells with Proguanil and Sulfasalazine illustrated strong anti-SARS-CoV-2 effects (represented by reductions of the envelope and nucleocapsid gene RNAs) in a dose dependent manner, mirroring the results of the initial screen (Figures 4B-E, S7B-E). Importantly, no significant effect on cellular viability was observed at any tested dose (Figures S7F-H). The effective concentration of sulfasalazine is comparable to maximal plasma concentrations achieved routinely in patients with rheumatoid arthritis or inflammatory bowel disease(IARC working Group on the Evaluation of Carcinogenic Risk to Humans, 2016).

**Figure 4.**
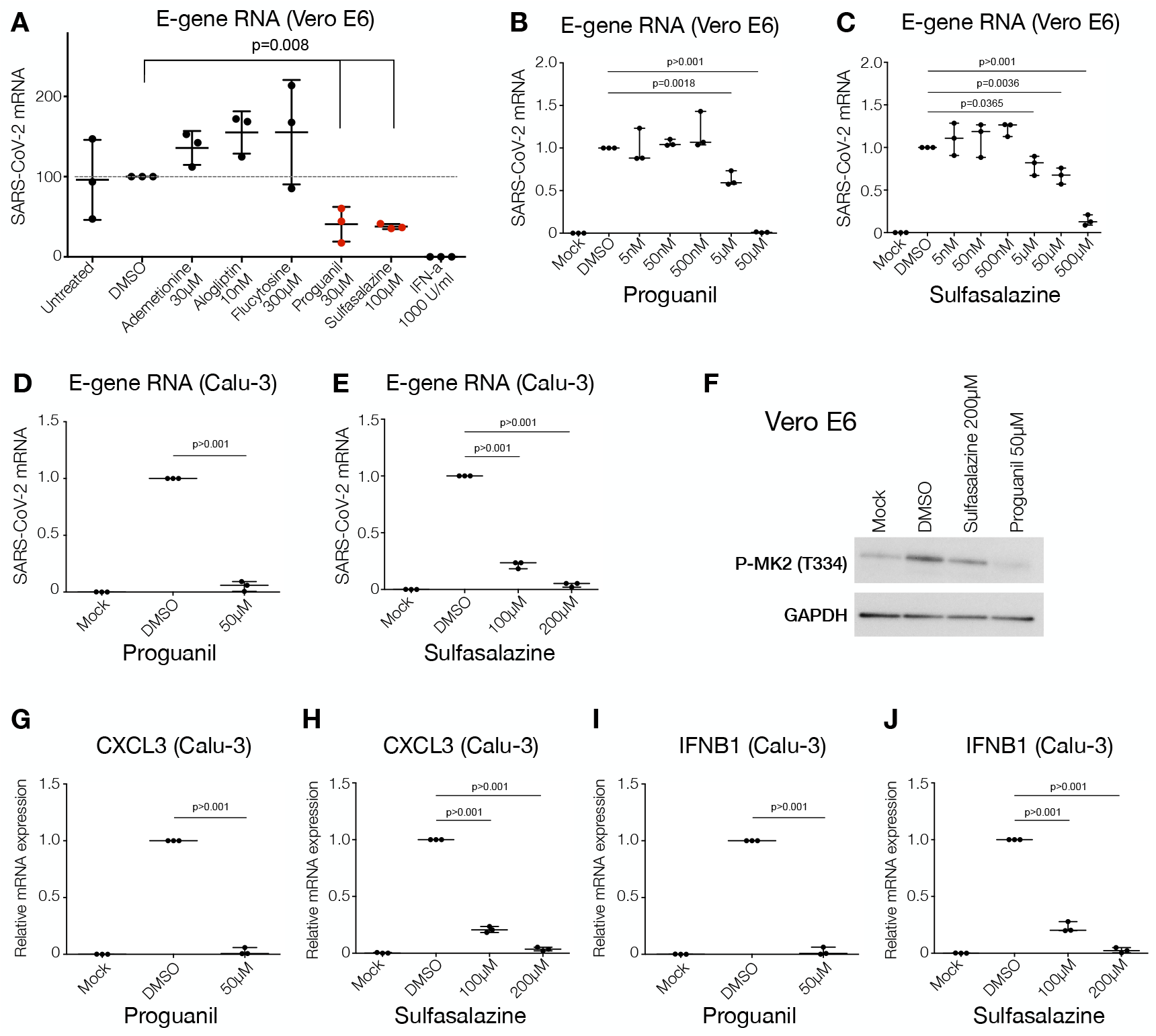
Proguanil and Sulfasalazine reduce SARS-CoV-2 replication and the p38/MAPK Signalling Activity. (A) qRT-PCR analysis of the indicated mRNA (Envelope, E-protein) from Vero E6 cells pre-treated with the indicated drugs and concentrations for three hours prior to infection with SARS-CoV-2 for 24 hours. Statistical test: Student’s t test. Mean + S.D. of three independent replicates is shown. (B, C) RT-qPCR analysis of indicated mRNA (Envelope, E-protein) from Vero E6 cells pre-treated with Proguanil or Sulfasalazine at indicated concentrations for three hours prior to infection with SARS-CoV-2 for 24 hours. Statistical test: Student’s t test. Mean + S.D. of three independent replicates is shown. (D, E) RT-qPCR analysis of indicated mRNA (Envelope, E-protein) from Calu-3 cells pre-treated with Proguanil or Sulfasalazine at indicated concentrations for three hours prior to infection with SARS-CoV-2 for 24 hours. Statistical test: Student’s t test. Mean + S.D. of three independent replicates is shown. (F) Western blot analysis of phosphorylated MAPKAPK-2 (Thr334) in Mock, DMSO, Sulfasalazine or Proguanil-treated Vero E6 cells at indicated concentrations for three hours prior to infection with SARS-CoV-2 for 24 hours. (G-J) RT-qPCR analysis of the indicated mRNAs from Calu-3 cells pre-treated with Proguanil or Sulfasalazine at indicated concentrations for three hours prior to infection with SARS-CoV-2 for 24 hours. Statistical test: Student’s t test. Mean + S.D. of three independent replicates is shown.

To further investigate the anti-SARS-CoV-2 impact of these two drugs, we examined the status of recently discovered intracellular pathways directly associated with SARS-CoV-2 infection and cytokine production(Bouhaddou et al., 2020). Indeed, treatment with either Proguanil or Sulfasalazine significantly reduced the phosphorylation of MAPKAPK2 (p-MK2, T334) (Figure 4F), an important component of the p38/mitogen-activated protein kinase (MAPK) signalling pathway, which has been shown to be activated via SARS-CoV-2 infection and stimulate cytokine response(Bouhaddou et al., 2020). Importantly, treatment of Calu-3 and Vero E6 cell lines with Proguanil and Sulfasalazine led to a significant downregulation of the mRNA of key cytokines (Figures 4G-J and S8), which are dictated by the p38/MAPK signalling pathway and shown to become elevated during SARS-CoV-2 infection and replication (*CXCL3*, *IFNB1* and *TNF-A*). Hence, the above results solidify the promising anti-SARS-CoV-2 effects of the two drugs, both at the viral as well as the molecular level.

To understand why Sulfasalazine and Proguanil are effective against SARS-CoV-2 infection, but others functioning in the same pathway were not (Figure 4A), we looked more closely at the targets of each drug. Figure 5 shows that SARS-CoV-2 orf8 binds to gamma-glutamyl hydrolase (GGH) and regulates the synthesis of NO, which is necessary for viral synthesis. An additional auxiliary pathway, mediating the synthesis of NADP, can also affect NO production, although indirectly. Sulfasalazine and Proguanil impinge on both of these pathways: Sulfasalazine targets the NFKB inhibitors NFKBIA and IKBKB as well as CYP450 enzymes, whereas Proguanil targets DHFR and CYP450 enzymes plus interacting partners. In this way these two drugs might more effectively target NO production and thus disrupt viral replication. By contrast, the three drugs that were not effective against SARS-CoV-2 infection (Flucytosine, Alogliptin and Ademetionine) only affect one of the two pathways. This analysis thereby highlights the possibility that targeting NO production through multiple pathways may be the reason for the efficacy of Sulfasalazine and Proguanil in reducing viral replication.

**Figure 5.**
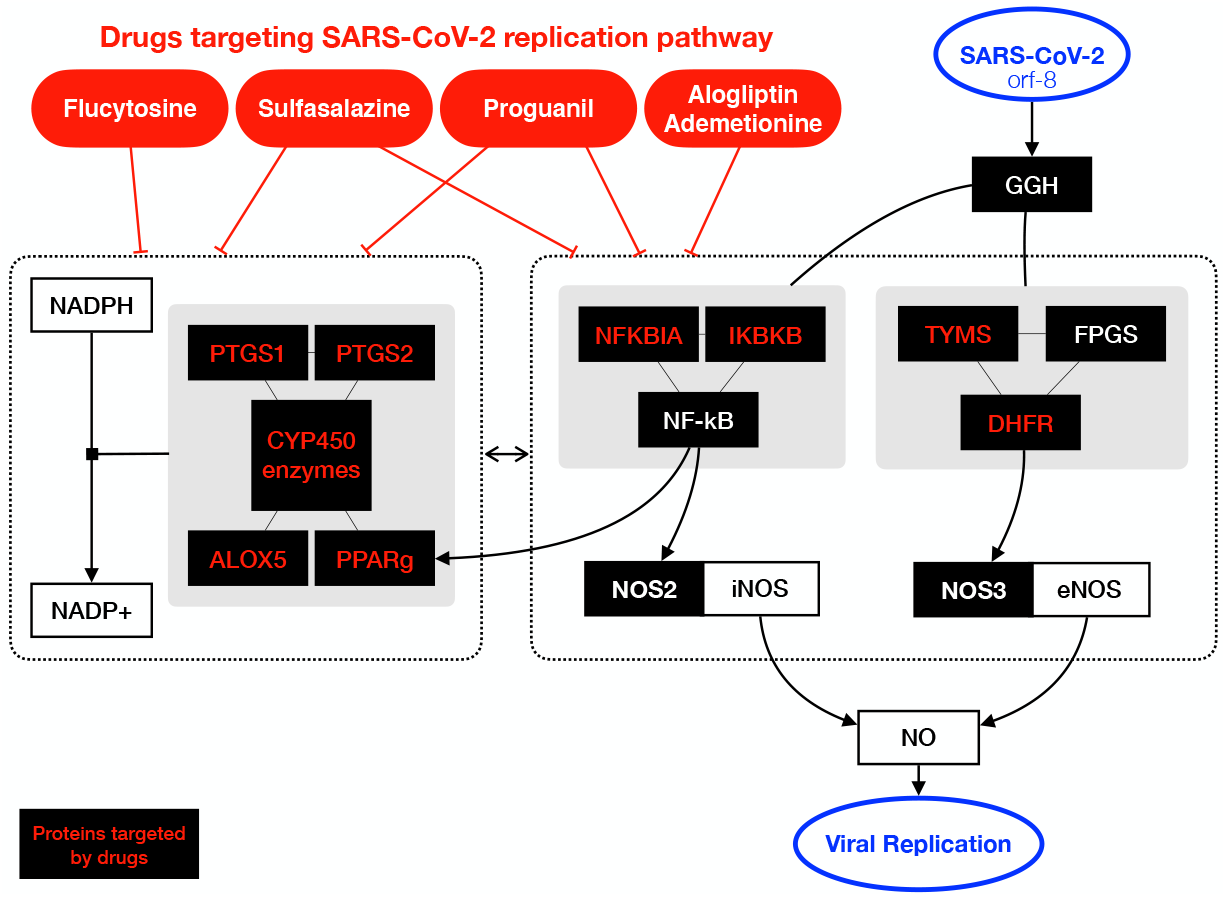
A schematics depicting the pathways mediating NO production that are targeted by the five tested drugs. The black boxes indicate key proteins in SIP network, and those targeted by the five drugs are highlighted in red colour. Sulfasalazine and Proguanil target proteins in both pathways that directly and indirectly (via NADP production) affect NO production (Choi et al., 2017; Corpas and Barroso, 2014; Hiscott et al., 2001; Wink et al., 2011).

## DISCUSSION

Here we have used a series of computational approaches, including bespoke methods for data integration, network analysis, computer simulation and machine learning, to identify novel SARS-CoV-2 induced pathways that could be targeted therapeutically by repurposing existing and approved drugs (Figure S9). Although network analysis is increasingly being used for analysis of genetic datasets to uncover disease signatures(Barabási et al., 2011), a few key aspects of our approach were essential in uncovering these new targets, including agnostic construction of the SIP network and application of novel algorithms (previously used in other industries including social media). In addition, the use of Artificial Neural Networks to understand systematically the mechanism of action for the drugs was vital to this investigation.

Our analysis identifies 200 approved drugs, along with their MoA, that may effective against COVID-19 (Table S2). We are confident that these drugs have a potential for repurposing for COVID-19, since 40 out of the 200 drugs have already entered clinical trials, testifying to the predictive power of our approach. An important part of our analysis is the use of already approved drugs. This allows for the rapid advancement of the most promising of the 160 drugs which are not already in clinical trials.

We identify two drugs, Sulfasalazine and Proguanil, that can reduce SARS-CoV-2 viral replication in cellular assays, raising the exciting possibility of their potential use in prophylaxis or treatment against COVID-19. Both of these drugs function through the NO pathway and have the potential to target more than one pathway necessary for NO production.

Safety is a particularly important consideration, since such drugs will be prescribed to any COVID-19 positive-case individuals who may have a broader range of underlying medical conditions and may not be hospitalised at the time of taking the drug. Sulfasalazine and Proguanil have the potential to be used prophylactically or therapeutically. Both drugs are well established and well tolerated drugs(Nakato et al., 2007; Nikfar et al., 2009). Sulfasalazine is already in use as an anti-inflammatory drug against autoimmune disorders. Given this drug has anti-viral activity (Figure 4), this raises the possibility that Sulfasalazine may act as an anti-viral and also an anti-inflammatory, if used against COVID-19. Proguanil is in used against malaria in combination with Atovaquone. It has an excellent safety profile and is well tolerated when used as a prophylactic and in treatment.

A complementary study to ours, that uses large-scale compound screening in cultured cells, has recently uncovered 100 molecules which have a partial effect on viral infectivity, 21 of which show a dose dependent reduction of viral replication(Riva et al., 2020). This list of drugs does not overlap with ours, with only two of our 200 approved drugs being present in this list. Neither Sulfasalazine nor Proguanil are amongst them. The main reason for this apparent disparity is that only 10% of the 100 compounds tested by Riva *et al.* are approved, whereas 100% of our 200 drugs are approved. This highlights the major difference in the two studies: our *in-silico* studies identify potential anti-viral drugs that are already approved and therefore at an advanced stage of repurposing, whereas Riva *et al.* have identified compounds validated in African Green Monkey cells, most of which are either in pre-clinical or phase 1-3 clinical trials.

Our study has shed unanticipated new light on COVID-19 disease mechanisms and has generated promising drug repurposing opportunities for prophylaxis and treatment. Our data-driven unsupervised approach and biological validation has uncovered 160 approved drugs not currently in clinical trials, which can be investigated immediately for repurposing and two drugs that show promise as anti-viral drugs. We expect this resource of potential drugs will facilitate and accelerate the development of therapeutics against COVID-19.

## Supporting information

SuppTable2

## ACKNOWLEDGMENTS

N.H. is funded by LifeArc. F.W. is funded by the LOEWE Center DRUID (project A3), the RAPID consortium of the Bundesministerium für Bildung und Forschung (BMBF, grant number 01KI1723E), and by the European Union’s Horizon 2020 research and innovation programme (MAD-CoV-2). K.T. is funded by a Wellcome Trust Sir Henry Wellcome Fellowship (grant reference RG94424). R.H. is funded by Cancer Research UK.

## AUTHOR CONTRIBUTIONS

N.H. and W.H. designed and performed the computational analyses. P.S. K.T. and E.Y. performed the experimental validation. N.H., R.H., K.T., F.W. and T.K. designed the experiments. M.M., W.L., N.M.K. and A.L. collected and analysed drug data. N.H., W.H., A.S., R.H., K.C., K.T. and T.K. interpreted results. F.M. reviewed drug list. N.H., K.T., F.W. and T.K. devised and supervised the project. N.H., W.H., A.S., K.T. and T.K. wrote the manuscript with contributions from all authors.

## DECLARATION OF INTERETES

A.L., A.S. and K.C. are funded by pharmaceutical industry. M.M. is an employee of LifeArc. E.Y. is funded by Storm Therapeutics. TK is a founder of Abcam and Storm Therapeutics.

## RESOURCE AVAILABILITY

### Code Availability

Computer programming scripts that were used in this study are available from https://github.com/wchwang/COVID19.

## Methods

### Directly Interacting Proteins (DIP) and Differentially Expressed Proteins (DEP)

332 high-confidence SARS-CoV-2-human PPIs(Gordon et al., 2020) were used as DIP. LC-MS/MS data at 6 hours, 24 hours after SARS-CoV-2 infection and no infection as a control(Bojkova et al., 2020) were analysed to identify Differentially Expressed Proteins (DEP) (|log2FC| > 1.5, FDR-BH p_adj_-value < 0.05).

### SIP network construction

SIP network was constructed of all shortest paths between DIP and DEP in a human protein-protein interaction network from STRING database(Szklarczyk et al., 2019). Only interactions with a confidence score of greater more than medium (0.4) were used. All shortest paths between DIP and DEP were found using the python package NetworkX(Hagberg et al., 2008). Networks were visualized using Gephi 0.9.2(Bastian et al., 2009) (Figure S1).

### Network analysis

Multiple network centrality algorithms were deployed to identify key proteins in SIP networks. Eigenvector centrality was used to identify the most influential proteins in the network. Degree centrality was used to identify the hub proteins in the network. Betweenness centrality was used to identify the bottleneck proteins in the network. Random walk with restart was used to identify proteins which are influenced by SARS-CoV-2. The algorithms were implemented in the python package NetworkX(Hagberg et al., 2008). Permutation tests were performed 1,000 times to identify significant proteins for each of the network centrality algorithms. For each permutation test, a random network that has the same degree distribution as the SIP network was generated. If a protein has less than permutation p-value 0.01 for each of the network centrality algorithms, we considered it a key protein.

### Key proteins functional enrichment analysis

Key proteins of SIP network were tested for enrichment of Jensen Disease(Pletscher-Frankild et al., 2015) and Gene Ontology (GO Biological Process) terms. Enrichment analyses were performed using Enrichr(Kuleshov et al., 2016).

### Visualization of a key network of SIP network

Key networks were built using interactions between the key proteins of the SIP network at 6 hours and 24 hours after infection. When visualising the key networks, subcellular localization of key proteins and enriched pathways of hidden layer proteins was added (Figures 2B and 2C). Subcellular localization information for key proteins was found using COMPARTMENT database(Binder et al., 2014). Among the available datasets in the COMPARTMENT database, ‘Knowledge channel’ data with a confidence score of greater than four was used. To identify enriched functions of the hidden layer proteins, the hidden layer proteins were tested for enrichment of Reactome pathway terms. Most hidden layer proteins belonged to the pathways “Metabolism of RNA”, “Cell cycle” and “Immune System” so we retained only these pathways for key network visualisation. The key networks visualization was carried out using Circos(Krzywinski et al., 2009).

### Drug-Target interactions

Approved drugs were collected from ChEMBL(Mendez et al., 2019) and DrugBank(Wishart et al., 2018). Drug-target interaction information was collected from DrugBank(v 5.1)(Wishart et al., 2018), STITCH(confidence score > 0.9)(Szklarczyk et al., 2016) and Cheng, et al(Cheng et al., 2019).

### *in-silico* drug efficacy simulation

Network-based *in-silico* drug efficacy simulation was conducted for key proteins from the SIP network at 6-hour and 24-hour. Given K, the set of key proteins from SIP networks, and T, the set of drug targets, the network proximity(equation (1)) of K with the target set of T of each approved drug where d(k, t) the shortest path length between nodes k ∈ K and t ∈ T in the human PPIs(Cheng et al., 2019) was executed.

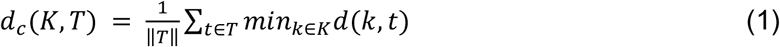

To assess the significance of the distance between a key protein of SIP network and a drug *d*_*c*_(*K,T*), the distance was converted to Z-score based on permutation tests by using

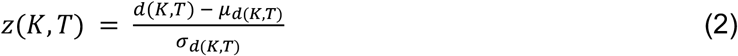

the permutation tests were repeated 1000 times, each time with two randomly selected gene lists with similar degree distributions to those of K and T. The corresponding p-value was calculated based on the permutation test results. Drug to SARS-CoV-2 associations with a Z-score of less than −2 were considered significantly proximal(Cheng et al., 2019).

### Drug-Pathway associations

Biological pathways were collated for the 200 drugs we identified by *in-silico* drug efficacy simulation. To do this, proteins targeted by the 200 drugs in SIP network were tested for enrichment of Reactome pathway terms using g:Profiler(Raudvere et al., 2019). Since Reacome pathway hierarchy contains main overarching parent pathways and more specific child pathways nested within these, in cases where child pathways were among enriched pathways, the parent pathway term was removed from the enriched pathways list. Finally, 148 Reactome pathways for 200 drugs were identified. Based on these identifications, a matrix containing 200 drugs and 148 Reactome pathways was generated for drug-pathway association. This matrix was constructed using the F1 score (F1=2(precision × recall)/(precision + recall) from the pathway enrichment analysis.

### MoA analysis

Self-Organizing Map (SOM)(KOHONEN, 1990) was used to analyse MoA of the 200 drugs. The data used in training was the F1 score matrix for Drug-Pathway associations (148 pathways by 200 drugs). After the SOM training, Davies-Bouldin index (DBI)(Davies and Bouldin, 1979) was calculated based on the U-matrix to determine the best patterning among partitions (Figure 3A). K-means algorithm were then used in order to find the 9 pathway clusters (Figure 3B). The SOM component maps of 148 pathways were analysed based on the clustering result and mapped into two MoA categories based on the biological functions (Figure S5). The SOM model also labelled each neurons with the 200 drugs (Figures 3D and 3E). The SOM Toolbox package(Vatanen et al., 2015) for Matlab was used for this analysis.

### Protein targeted by the 200 drugs

The frequency of drug-protein targeting was counted. Permutation tests were then performed 100 times to identify the significance threshold for the frequency of drug-protein targeting (Figure S6A). For each permutation test, the 200 drugs among all the drugs which we used for the *in silico* drug efficacy simulation were randomly selected. Then, the number of drugs targeting the same protein was calculated for all of the randomly selected 200 drugs. The proteins frequently targeted in the SIP network than randomised network were then tested for enrichment of UniProt Keywords (Figure S6B).

### Cell Culture

Chlorocebus sabaeus (Green monkey) Vero E6 (Vero 76, clone E6, Vero E6, ATCC® CRL-1586) authenticated by ATCC and tested negative for mycoplasma contamination prior to commencement were maintained in a humidified atmosphere at 37°C with 5% CO_2_, in Dulbecco’s modified Eagle’s medium (DMEM) containing 10% (v/v) fetal bovine serum (FBS, Invitrogen). Calu-3 (ATCC® HTB-55) human lung cells tested negative for mycoplasma contamination prior to commencement were maintained in a humidified atmosphere at 37°C with 5% CO_2_ in Eagle’s Minimum Essential Medium containing 20% (v/v) FBS. Human cell lines employed were either not listed in the cross-contaminated or misidentified cell lines database curated by the International Cell Line Authentication Committee (ICLAC) or were previously verified by karyotyping.

### Viruses and infections

Infection experiments were performed under biosafety level 3 conditions. SARS-CoV-2 (strain München-1.2/2020/984) isolate was propagated in Vero E6 cells in DMEM supplemented with 2% FBS. For infection experiments in Vero E6 and Calu-3 cells, SARS-CoV-2 (strain München-1.2/2020/984) at MOI=0.01 pfu/cell for 24 hours. All work involving live SARS-CoV-2 was performed in the BSL-3 facility of the Institute for Virology, University of Giessen (Germany), and was approved according to the German Act of Genetic Engineering by the local authority.

### Cell infection and drug treatment

Vero E6 and Calu-3 cells were seeded using 8×10^4^ cells in 24-well plates. The following day cells were treated for 3 hour prior to infection with the indicated doses of Ademethionine (30μM, Selleckchem), Alogliptin (10μM, Selleckchem), Flucytosine (300μM, Selleckchem), Proguanil (5nM-500μM, Selleckchem), Sulfasalazine (5nM-500μM, Selleckchem), IFN-A (1000 U/ml), DMSO (Sigma) or mock and infected with SARS-CoV-2 at MOI of 0.01 in serum-free DMEM at 37°C for 24 hours before RNA or protein lysis. Infection experiments were performed under biosafety level 3 conditions.

### Quantitative RT-PCR analysis

RNA was isolated using the RNeasy Mini (Qiagen). SARS-CoV-2 replication (E-gene and N-gene RNA) and gene expression of the cytokines *CXCL3, IFNB1 and TNF-A* was quantified by RT-qPCR. For cDNA synthesis, RNA was reverse-transcribed with the SuperScript VILO cDNA Synthesis kit (Invitrogen, 11755-050). The levels of specific RNAs were measured using the ABI 7900 real-time PCR machine and the PowerUp™ SYBR™ Green Master Mix (Applied Biosystems, 100029284) according to the manufacturer’s instructions. ΔCT values were determined relative to the GAPDH and ΔΔCT values were normalized to infected DMSO treated samples. Error bars indicate the standard deviation of the mean from three independent biological replicates. All primer sequences are listed in Table 1 below.

**Table 1.**
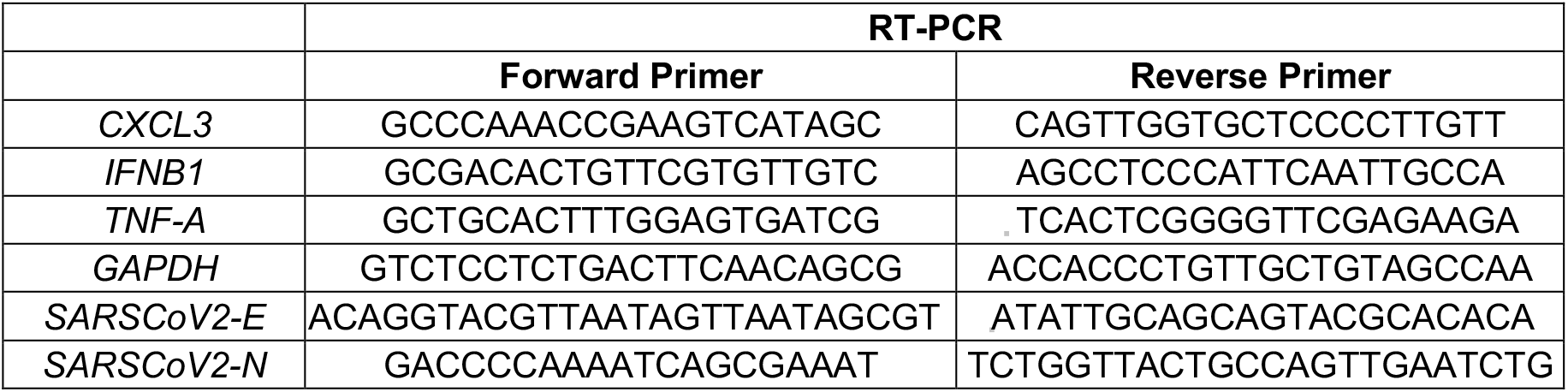

### Cytotoxicity cell viability assays

Cytotoxicity was performed in Vero E6 and Calu-3 cells using Neutral Red (Abcam, ab234039) and MTT assay (Roche) respectively, according to the manufacturer’s instructions. Cytotoxicity was performed in Vero E6 and Calu-3 cells with the indicated compound dilutions and concurrent with viral replication assays. All assays were performed in biologically independent triplicates.

### Western blot analysis

8×10^4^ Vero E6 cells either mock-infected or infected and treated with DMSO or Proguanil (50μM) or Sulfasalazine (200μM) for 24 hours, were resuspended and lysed in whole cell 1×SDS sample buffer (4×SDS sample buffer: 143 mM Tris-HCl, pH = 6.8, 28.6% Glycerol, 5.7% SDS, 4.3 mM Bromophenol Blue), supplemented with 2ml 2-mercaptoethanol, protease inhibitors (Sigma), and phosphatase inhibitors (Sigma) and boiled for 5 min at 95°C. 10-20 μg of protein was separated on SDS-PAGE gels, and blotted onto polyvinylidene difluoride membranes (Millipore).

### Antibodies

Western blot experiments were performed using the following antibodies: GAPDH (Abcam, ab9484), Phospho-MAPKAPK-2 (Thr334, Cell Signalling, 3007), Goat anti-Rabbit (Abcam, ab6721) and Anti-mouse-HRP (Cell Signalling, 7076S).

### Statistical analysis

Statistical analyses performed were specified in figure legends. Differences were considered significant for P-values < 0.05.

## REFERENCES

Barabási, A.L., Gulbahce, N., and Loscalzo, J. (2011). Network medicine: A network-based approach to human disease. Nat. Rev. Genet. 12, 56–68.

Bastian, M., Heymann, S., and Jacomy, M. (2009). Gephi: An open source software for exploring and manipulating networks. BT - International AAAI Conference on Weblogs and Social. Int. AAAI Conf. Weblogs Soc. Media 361–362.

Binder, J.X., Pletscher-Frankild, S., Tsafou, K., Stolte, C., O’Donoghue, S.I., Schneider, R., and Jensen, L.J. (2014). COMPARTMENTS: Unification and visualization of protein subcellular localization evidence. Database 2014, 1–9.

Blanco-Melo, D., Nilsson-Payant, B.E., Liu, W.-C., Uhl, S., Hoagland, D., Møller, R., Jordan, T.X., Oishi, K., Panis, M., David Sachs, et al. (2020). Imbalanced host response to SARS-CoV-2 drives development of COVID-19. Cell.

Bojkova, D., Klann, K., Koch, B., Widera, M., Krause, D., Ciesek, S., Cinatl, J., and Münch, C. (2020). Proteomics of SARS-CoV-2-infected host cells reveals therapy targets. Nature.

Bouhaddou, M., Memon, D., Meyer, B., White, K.M., Rezelj, V. V., Correa Marrero, M., Polacco, B.J., Melnyk, J.E., Ulferts, S., Kaake, R.M., et al. (2020). The Global Phosphorylation Landscape of SARS-CoV-2 Infection. Cell 1–28.

Cheng, F., Kovács, I.A., and Barabási, A.L. (2019). Network-based prediction of drug combinations. Nat. Commun.

Choi, M.J., Lee, E.J., Park, J.S., Kim, S.N., Park, E.M., and Kim, H.S. (2017). Anti-inflammatory mechanism of galangin in lipopolysaccharide-stimulated microglia: Critical role of PPAR-γ signaling pathway. Biochem. Pharmacol. 144, 120–131.

Clinicaltrials.gov (2020). SARS-CoV-2 Clinical trials.

Corpas, F.J., and Barroso, J.B. (2014). NADPH-generating dehydrogenases: Their role in the mechanism of protection against nitro-oxidative stress induced by adverse environmental conditions. Front. Environ. Sci. 2, 1–5.

Courtney J. Mycroft-West, Dunhao Su, Yong Li, Scott E. Guimond, Timothy R. Rudd, S., and Elli, Gavin Miller, Quentin M. Nunes, Patricia Procter, Antonella Bisio, Nicholas R. Forsyth, Jeremy E. Turnbull, Marco Guerrini, David G. Fernig, Edwin A. Yates, M.A.L. and M.A.S. (2020). Glycosaminoglycans induce conformational change in the SARS-. BioRxiv April 29.

Criado, P.R., Pagliari, C., Carneiro, F.R.O., and Quaresma, J.A.S. (2020). Lessons from dermatology about inflammatory responses in Covid-19. Rev. Med. Virol. 1–18.

Davies, D.L., and Bouldin, D.W. (1979). A Cluster Separation Measure. IEEE Trans. Pattern Anal. Mach. Intell. PAMI-1, 224–227.

Gordon, D.E., Jang, G.M., Bouhaddou, M., Xu, J., Obernier, K., White, K.M., O’meara, M.J., Rezelj, V. V, Guo, J.Z., Swaney, D.L., et al. (2020). A SARS-CoV-2 protein interaction map reveals targets for drug repurposing. Nature.

Guan, W., Ni, Z., Hu, Y., Liang, W., Ou, C., He, J., Liu, L., Shan, H., Lei, C., Hui, D.S.C., et al. (2020). Clinical Characteristics of Coronavirus Disease 2019 in China. N. Engl. J. Med. 382, 1708–1720.

Guney, E., Menche, J., Vidal, M., and Barábasi, A.-L. (2016). Network-based in silico drug efficacy screening. Nat. Commun. 7, 10331.

Gupta, A., Madhavan, M. V., Sehgal, K., Nair, N., Mahajan, S., Sehrawat, T.S., Bikdeli, B., Ahluwalia, N., Ausiello, J.C., Wan, E.Y., et al. (2020). Extrapulmonary manifestations of COVID-19. Nat. Med. 26.

Hagberg, A.A., Schult, D.A., and Swart, P.J. (2008). Exploring network structure, dynamics, and function using NetworkX. 7th Python Sci. Conf. (SciPy 2008) 11–15.

Hiscott, J., Kwon, H., and Génin, P. (2001). Hostile takeovers: Viral appropriation of the NF-κB pathway. J. Clin. Invest. 107, 143–151.

IARC working Group on the Evaluation of Carcinogenic Risk to Humans (2016). Some Drugs and Herbal Products.

Kohonen, T. (1990). The Self-organizing Map. Proc. IEEE 78, 1464–1480.

Krzywinski, M., Schein, J., Birol, I., Connors, J., Gascoyne, R., Horsman, D., Jones, S.J., and Marra, M.A. (2009). Circos: An information aesthetic for comparative genomics. Genome Res. 19, 1639–1645.

Kuleshov, M., Clarke, D.J.B., Kropiwnicki, E., Jagodnik, K., Bartal, A., Evangelista, J.E., Zhou, A., Ferguson, L.B., Lachmann, A., and Ma’ayan, A. (2020). The COVID-19 Gene and Drug Set Library. SSRN Electron. J.

Kuleshov, M. V., Jones, M.R., Rouillard, A.D., Fernandez, N.F., Duan, Q., Wang, Z., Koplev, S., Jenkins, S.L., Jagodnik, K.M., Lachmann, A., et al. (2016). Enrichr: a comprehensive gene set enrichment analysis web server 2016 update. Nucleic Acids Res. 44, W90–W97.

kumar, arun; C.S, Sharanya; J, Abhithaj; c, S. (2020). Drug Repurposing to Identify Therapeutics Against COVID 19 with SARS-Cov-2 Spike Glycoprotein and Main Protease as Targets: An in Silico Study. ChemRxiv.

Kwiecien, S., Jasnos, K., Magierowski, M., Sliwowski, Z., Pajdo, R., Brzozowski, B., Mach, T., Wojcik, D., and Brzozowski, T. (2014). Lipid peroxidation, reactive oxygen species and antioxidative factors in the pathogenesis of gastric mucosal lesions and mechanism of protection against oxidative stress - induced gastric injury. J. Physiol. Pharmacol. 65, 613–622.

Lind, M., Hayes, A., Caprnda, M., Petrovic, D., Rodrigo, L., Kruzliak, P., and Zulli, A. (2017). Inducible nitric oxide synthase: Good or bad? Biomed. Pharmacother. 93, 370–375.

Mendez, D., Gaulton, A., Bento, A.P., Chambers, J., De Veij, M., Félix, E., Magariños, M.P., Mosquera, J.F., Mutowo, P., Nowotka, M., et al. (2019). ChEMBL: Towards direct deposition of bioassay data. Nucleic Acids Res. 47, D930–D940.

MirIbrahim Sajid, Javeria Tariq, SheharBano Awais, ZehraNaseem, SamiraShabbir Balouch, S.A. (2020). SARS-CoV-2: Cytokine Storm and Therapy. Ann. King Edward Med. Univ. 26, 243–251.

Nakato, H., Vivancos, R., and Hunter, P.R. (2007). A systematic review and meta-analysis of the effectiveness and safety of atovaquone - Proguanil (Malarone) for chemoprophylaxis against malaria. J. Antimicrob. Chemother. 60, 929–936.

Nikfar, S., Rahimi, R., Rezaie, A., and Abdollahi, M. (2009). A meta-analysis of the efficacy of sulfasalazine in comparison with 5-aminosalicylates in the induction of improvement and maintenance of remission in patients with ulcerative colitis. Dig. Dis. Sci. 54, 1157–1170.

Pletscher-Frankild, S., Pallejà, A., Tsafou, K., Binder, J.X., and Jensen, L.J. (2015). DISEASES: Text mining and data integration of disease-gene associations. Methods 74, 83–89.

Raudvere, U., Kolberg, L., Kuzmin, I., Arak, T., Adler, P., Peterson, H., and Vilo, J. (2019). G:Profiler: A web server for functional enrichment analysis and conversions of gene lists (2019 update). Nucleic Acids Res. 47, W191–W198.

Riva, L., Yuan, S., Yin, X., Martin-Sancho, L., Matsunaga, N., Pache, L., Burgstaller-Muehlbacher, S., De Jesus, P.D., Teriete, P., Hull, M. V, et al. (2020). Discovery of SARS-CoV-2 antiviral drugs through large-scale compound repurposing. Nature.

Romero-Brey, I., and Bartenschlager, R. (2016). Endoplasmic reticulum: The favorite intracellular niche for viral replication and assembly. Viruses 8, 1–26.

Shin, D., Mukherjee, R., Grewe, D., Bojkova, D., Baek, K., Bhattacharya, A., Schulz, L., Widera, M., Mehdipour, A.R., Tascher, G., et al. (2020). Papain-like protease regulates SARS-CoV-2 viral spread and innate immunity. Nature.

Szklarczyk, D., Santos, A., von Mering, C., Jensen, L.J., Bork, P., and Kuhn, M. (2016). STITCH 5: augmenting protein– chemical interaction networks with tissue and affinity data. Nucleic Acids Res. 44, D380–D384.

Szklarczyk, D., Gable, A.L., Lyon, D., Junge, A., Wyder, S., Huerta-Cepas, J., Simonovic, M., Doncheva, N.T., Morris, J.H., Bork, P., et al. (2019). STRING v11: Protein-protein association networks with increased coverage, supporting functional discovery in genome-wide experimental datasets. Nucleic Acids Res. 47, D607–D613.

Tang, X., Wu, C., Li, X., Song, Y., Yao, X., Wu, X., Duan, Y., Zhang, H., Wang, Y., Qian, Z., et al. (2020). On the origin and continuing evolution of SARS-CoV-2. Natl. Sci. Rev. 7, 1012–1023.

Vatanen, T., Osmala, M., Raiko, T., Lagus, K., Sysi-Aho, M., Orešič, M., Honkela, T., and Lähdesmäki, H. (2015). Self-organization and missing values in SOM and GTM. Neurocomputing 147, 60–70.

Vesanto, J., and Alhoniemi, E. (2000). Clustering of the Self−Organizing Map. IEEE Trans. Neural Networks 11, 586–600.

Wang, G., Moniri, N.H., Ozawa, K., Stamler, J.S., and Daaka, Y. (2006). Nitric oxide regulates endocytosis by S-nitrosylation of dynamin. Proc. Natl. Acad. Sci. U. S. A. 103, 1295–1300.

Wink, D.A., Hines, H.B., Cheng, R.Y.S., Switzer, C.H., Flores-Santana, W., Vitek, M.P., Ridnour, L.A., and Colton, C.A. (2011). Nitric oxide and redox mechanisms in the immune response. J. Leukoc. Biol. 89, 873–891.

Wishart, D.S., Feunang, Y.D., Guo, A.C., Lo, E.J., Marcu, A., Grant, J.R., Sajed, T., Johnson, D., Li, C., Sayeeda, Z., et al. (2018). DrugBank 5.0: a major update to the DrugBank database for 2018. Nucleic Acids Res. 46, D1074–D1082.

