## Supplemental Figures and Tables for "Identification of SARS-CoV-2 induced pathways reveal drug repurposing strategies"

Figure S1

A

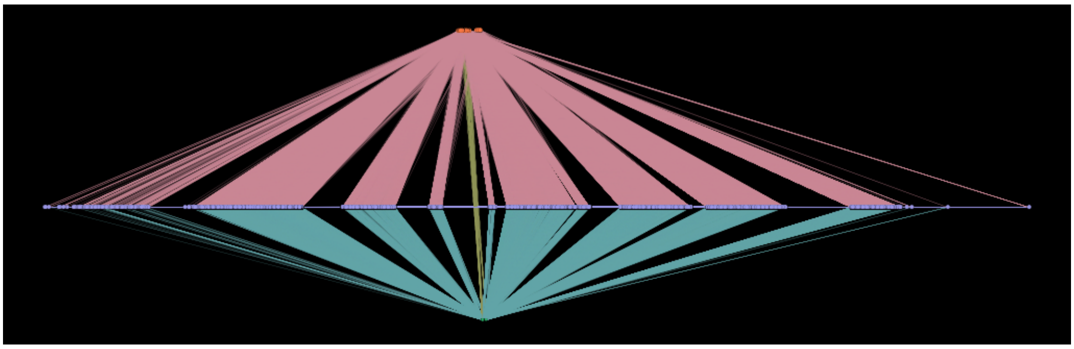

B

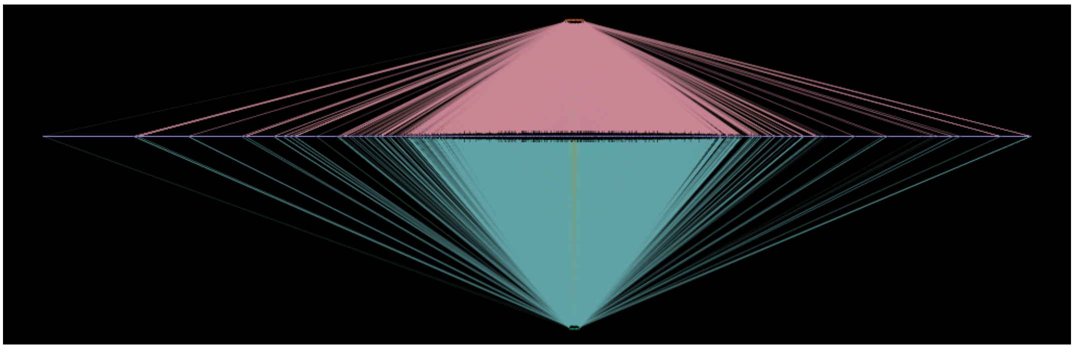

**Figure S1** SARS-CoV-2-induced protein (SIP) network (A) 6-hour network (B) 24-hour network

Figure S2

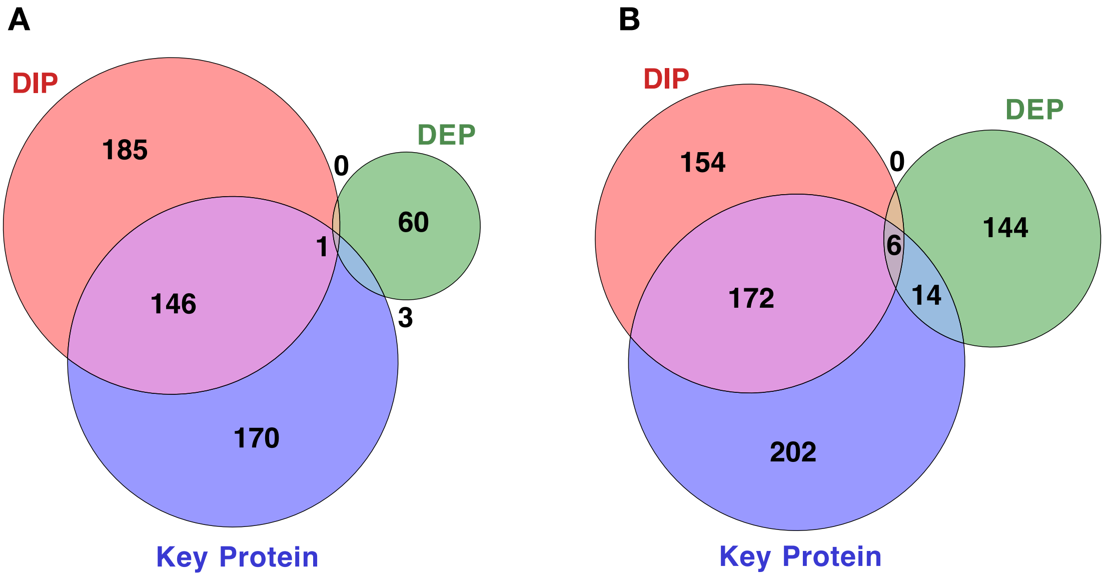


**Figure S2** Venn diagrams (A) DEP, DIP and 6-hour key proteins (B) DEP, DIP and 24-hour key proteins

Figure S3

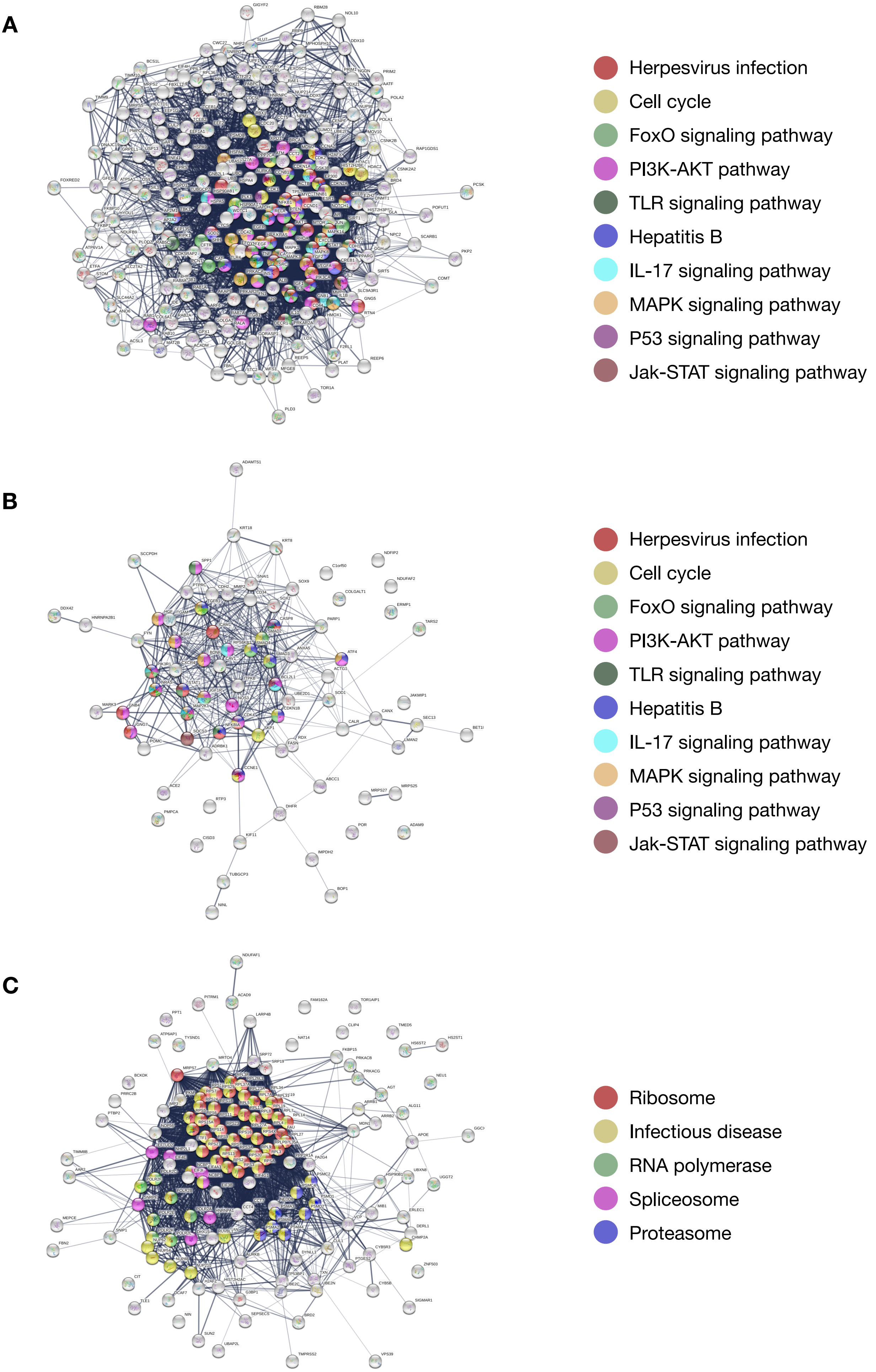


**Figure S3** SIP network for key proteins (A) 6-hour and 24-hour common, (B) 6-hour specific, (C) 24-hour specific

Figure S4

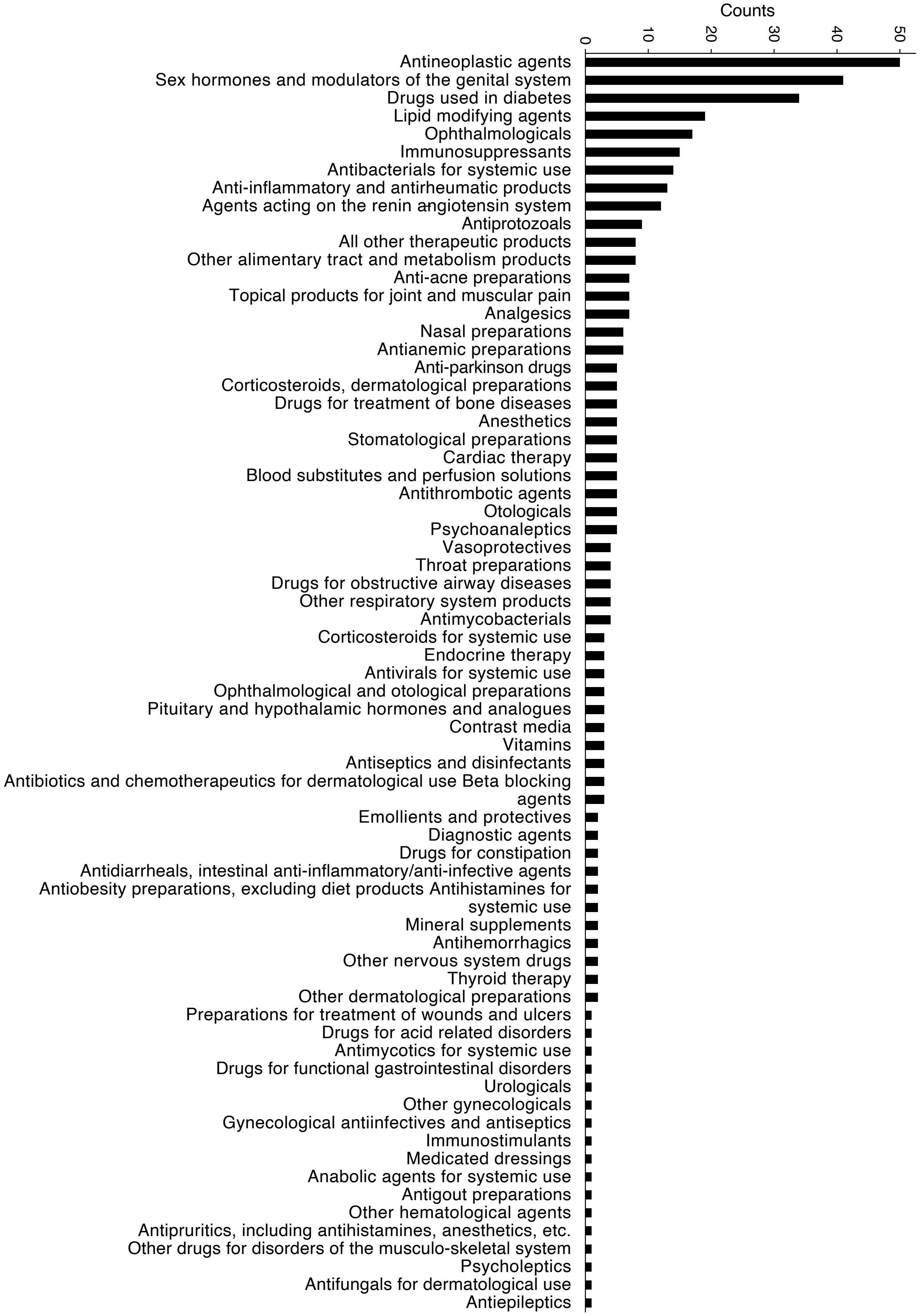


**Figure S4** Enriched ATC codes for the 240 identified compounds

Figure S5

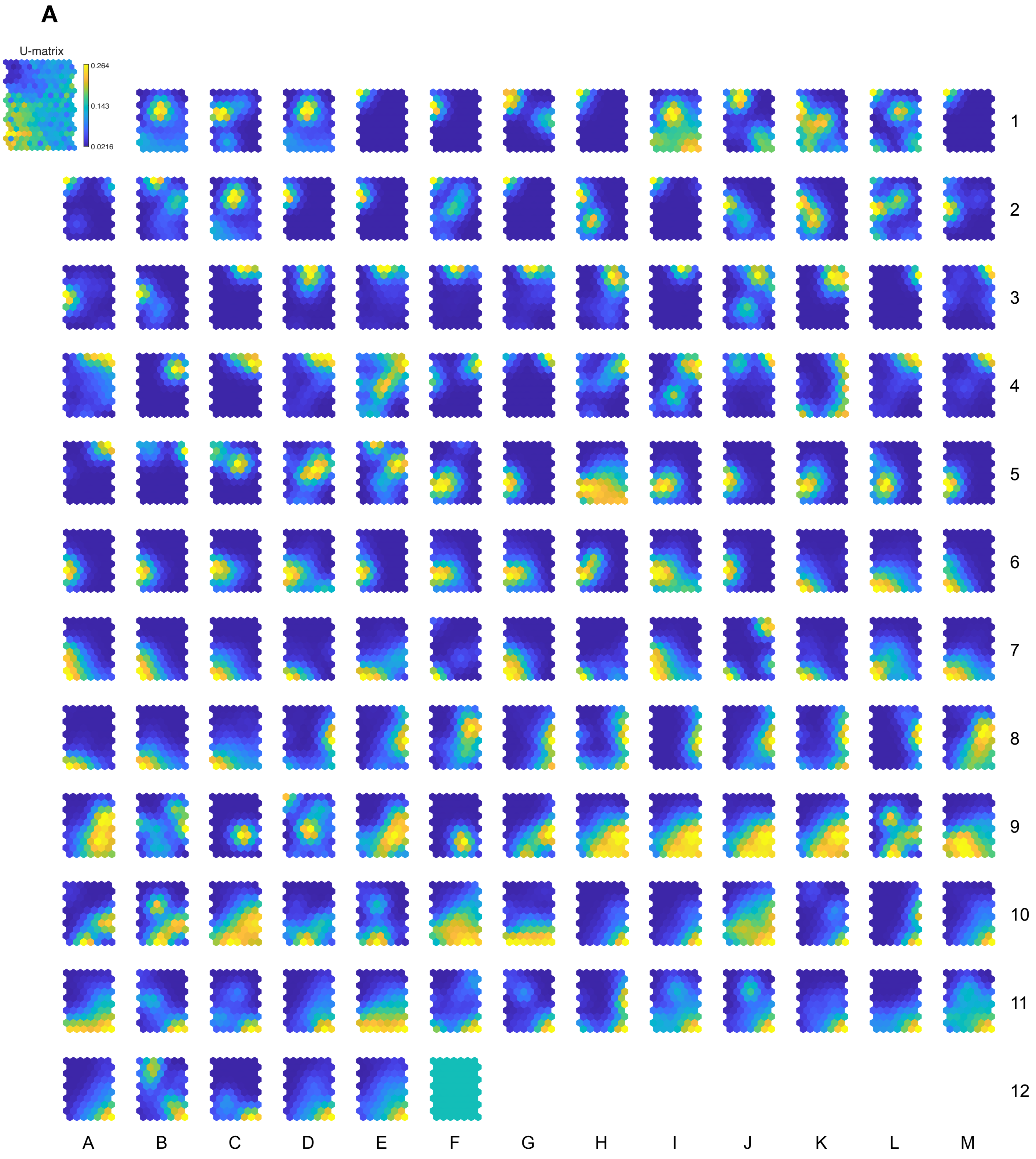


**Figure S5** SOM results of 148 pathways. (A) SOM visualization of U-matrix (top left) and 148 component maps, one for each pathway. The component layers are linked following similarity colour and position. The 148 figures are linked by position: in each map, the hexagon in a certain position corresponds to the same map unit. The component planes show which pathways have similar target-pathway association. The U-matrix shows the distances between the neighbouring neurons hence visualizes the cluster structure of the map. For instance, lower values (dark blue) on the U-matrix indicate a shorter distance between neighbouring neurons hence the cluster can be formed. (B) The name of the component planes.

Figure S5

## B

| A | B | C | D | E | F | G | H | I | J | K | L | M | |
|---|---|---|---|---|---|---|---|---|---|---|---|---|---|
| | Apoptotic factor-mediated response | Cooperation of Prefoldin and TriC/CCT in actin and tubulin folding | Cytochrome c-mediated apoptotic response | Defects in cobalamin (B12) metabolism | Digestion | Incretin synthesis, secretion, and inactivation | Intestinal absorption | Intrinsic Pathway for Apoptosis | Loss of Function of TGFBR1 in Cancer | Metabolism of carbohydrat es | Metabolism of nitric oxide: eNOS activation and regulation | mitochondria l fatty acid beta-oxidation of saturated fatty acids | 1 |
| Organic cation/ anion/ zwitterion transport | Plasma lipoprotein assembly | SMAC, XIAP-regulated apoptotic response | Sphingolipid metabolism | Synthesis of substrates in N-glycan biosysthesis | The citric acid (TCA) cycle and respiratory electron transport | Thyroxine biosynthesis | Transport of Mature mRNAs Derived from Intronless Transcripts | Abacavir transport and metabolism | Cap-dependent Translation Initiation | Influenza Infection | Metabolism of cofactors | Metabolism of water-soluble vitamins and cofactors | 2 |
| Nucleotide salvage | Selenoamin o acid metabolism | Acetylcholin e binding and downstream events | Amine Oxidase reactions | Biosynthesis of DHA- derived SPMs | Biosynthesis of maresins | Biosynthesis of specialized proresolving mediators (SPMs) | Gamma-carboxylatio n, transport, and amino-terminal cleavage of proteins | Presynaptic nicotinic acetylcholin e receptors | Regulation of cholesterol biosynthesis by SREBP (SREBF) | Synthesis of bile acids and bile salts | Transport of vitamins, nucleosides, and related molecules | Amine ligand-binding receptors | 3 |
| Arachidonic acid metabolism | Binding and Uptake of Ligands by Scavenger Receptors | Cardiac conduction | Cytochrome P450 - arranged by substrate type | Fatty acid metabolism | Glycerophos pholipid biosynthesis | Metabolism of amine-derived hormones | Metabolism of porphyrins | Metabolism of steroids | Neurotrans mitter clearance | Neurotrans mitter receptors and postsynaptic signal transmission | Phase I - Functionaliz ation of compounds | Phase II - Conjugation of compounds | 4 |
| Serotonin clearance from the synaptic cleft | Transport of bile salts and organic acids, metal ions and amine compounds | Nitric oxide stimulates guanylate cyclase | Response to metal ions | Plasma lipoprotein remodeling | Activation of HOX genes during differentiatio n | Base Excision Repair | Cellular Senescence | Chromatin modifying enzymes | DNA strand elongation | Epigenetic regulation of gene expression | Gene Silencing by RNA | Global Genome Nucleotide Excision Repair (GG-NER) | 5 |
| HDR through Homologous Recombinati on (HRR) | HDR through Homologous Recombinati on (HRR) or Single Strand Annealing | Mitotic Prometapha se | Regulation of TP53 Activity | Resolution of D-Loop Structures | SUMO E3 ligases SUMOylate target proteins | SUMOylatio n | Transcriptio n of the HIV genome | Transcriptio nal Regulation by TP53 | Translesion synthesis by Y family DNA polymerases bypasses lesions on DNA | APC/C-mediated degradation of cell cycle proteins | Deubiquitina tion | DNA Replication Pre-Initiation | 6 |
| G1/S Transition | G2/M Checkpoints | G2/M Transition | Hedgehog 'off' state | Interleukin-1 family signaling | Ion channel transport | M Phase | Membrane Trafficking | Mitotic G1 phase and G1/S transition | Neurotrans mitter release cycle | Regulation of mRNA stability by proteins that bind AU-rich elements | Selective autophagy | Signaling by NOTCH | 7 |
| Signaling by the B Cell Receptor (BCR) | TCF dependent signaling in response to WNT | Transcriptio nal regulation by RUNX3 | Anti-inflammatory response favouring Leishmania parasite infection | Class A/1 (Rhodopsin-like receptors) | Formation of Fibrin Clot (Clotting Cascade) | G alpha (q) signalling events | G-protein mediated events | GPCR downstream signalling | GPCR ligand binding | Opioid Signalling | Potassium Channels | Regulation of Insulin-like Growth Factor (IGF) transport and uptake by Insulin-like Growth | 8 |
| Response to elevated platelet cytosolic $Ca^{2+}$ | Unfolded Protein Response (UPR) | Diseases of glycosylation | Iron uptake and transport | Degradation of the extracellular matrix | Diseases associated with glycosamino glycan metabolism | Diseases associated with the TLR signaling cascade | MyD88 dependent cascade initiated on endosome | Toll Like Receptor 2 (TLR2) Cascade | Toll Like Receptor 9 (TLR9) Cascade | Toll-like Receptor Cascades | Caspase activation via Death Receptors in the presence of ligand | Generic Transcriptio n Pathway | 9 |
| Interferon alpha/beta signaling | Regulated Necrosis | Signaling by Interleukins | TNF signaling | TP53 Regulates Transcriptio n of Cell Death Genes | FOXO-mediated transcription | Axon guidance | Downstream signaling of activated FGFR1 | Downstream signaling of activated FGFR3 | ESR-mediated signaling | FGFR2 ligand binding and activation | G-protein beta:gamma signalling | Insulin receptor signalling cascade | 10 |
| MAPK1/MA PK3 signaling | mTOR signalling | Negative regulation of MAPK pathway | Oncogenic MAPK signaling | PIP3 activates AKT signaling | Post NMDA receptor activation events | Prolonged ERK activation events | Regulation of insulin secretion | Signaling by ERBB2 | Signaling by ERBB2 in Cancer | Signaling by FGFR | Signaling by MET | Signaling by Non- Receptor Tyrosine Kinases | 11 |
| Signaling by NTRK2 (TRKB) | Signaling by TGF-beta Receptor Complex in Cancer | Signaling by WNT in cancer | Signalling to ERKs | VEGFA-VEGFR2 Pathway | Uptake and actions of bacterial toxins | | | | | | | | 12 |

Figure S6

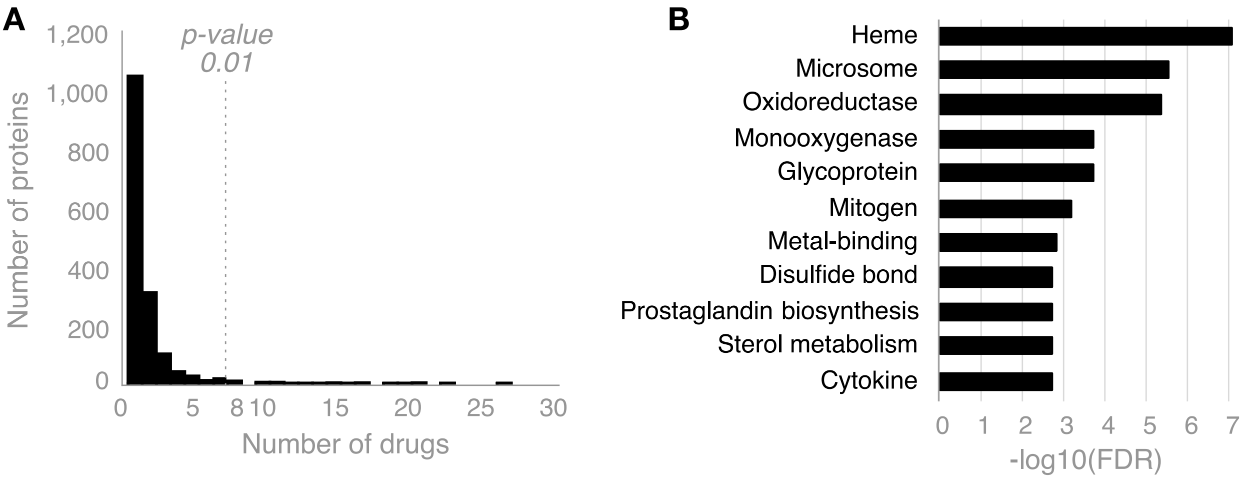


**Figure S6** (A) histogram of the number of drugs targeting specific proteins. A subset of 1,573 proteins in the SIP network are targeted by 200 drugs, and the number of drugs targeting a protein ranges from 1 to 27. The p-value of the permutation test is shown as a cut-off point. (B) The Uniprot Keyword enrichment test for the top 30 proteins that are hit by 8 or more drugs.

Figure S7

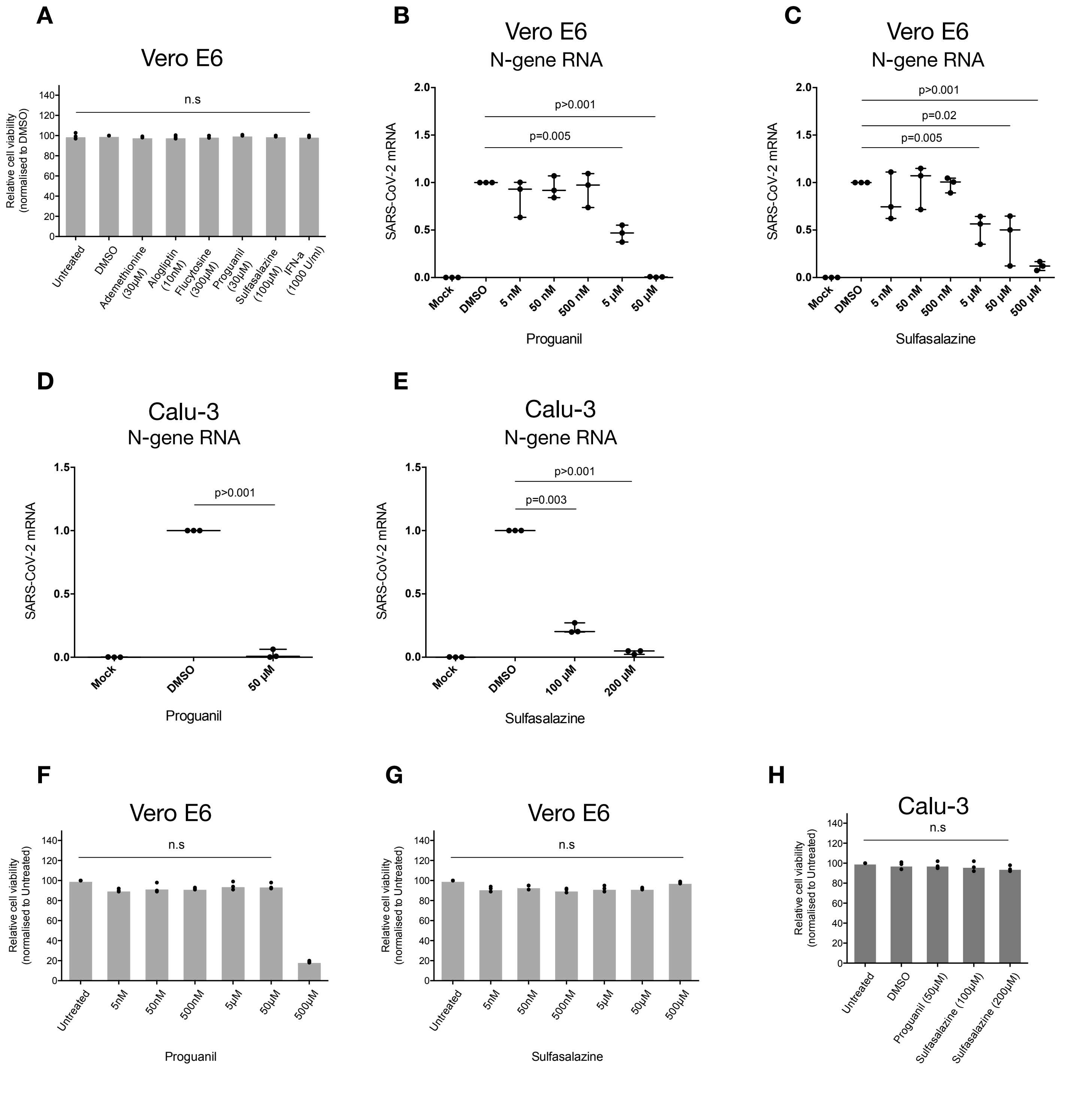


**Figure S7** Proguanil and Sulfasalazine significantly impair SARS-CoV-2 replication and blocked the p38/MAPK Signalling Activity. (A) Cell viability analysis of Vero E6 cells treated with the indicated drugs and concentrations for 24 hours. Statistical test: Student’s t test. Mean + S.D. of three independent replicates is shown. n.s., not significant. (B and C) RT-qPCR analysis of indicated mRNA (Nucleocapsid, N-protein) from Vero E6 cells pre-treated with Proguanil or Sulfasalazine at indicated concentrations for three hours prior to infection with SARS-CoV-2 for 24 hours. Statistical test: Student’s t test. Mean + S.D. of three independent replicates is shown. (D and E) RT-qPCR analysis of indicated mRNA (Nucleocapsid, N-protein) from Calu-3 cells pre-treated with Proguanil or Sulfasalazine at indicated concentrations for three hours prior to infection with SARS-CoV-2 for 24 hours. Statistical test: Student’s t test. Mean + S.D. of three independent replicates is shown. (F and G) Cell viability analysis of Vero E6 cells treated with the indicated drugs and concentrations for 24 hours. Statistical test: Student’s t test. Mean + S.D. of three independent replicates is shown. n.s., not significant. (H) Cell viability analysis of Calu-3 cells treated with the indicated drugs and concentrations for 24 hours. Statistical test: Student’s t test. Mean + S.D. of three independent replicates is shown. n.s., not significant.

Figure S8

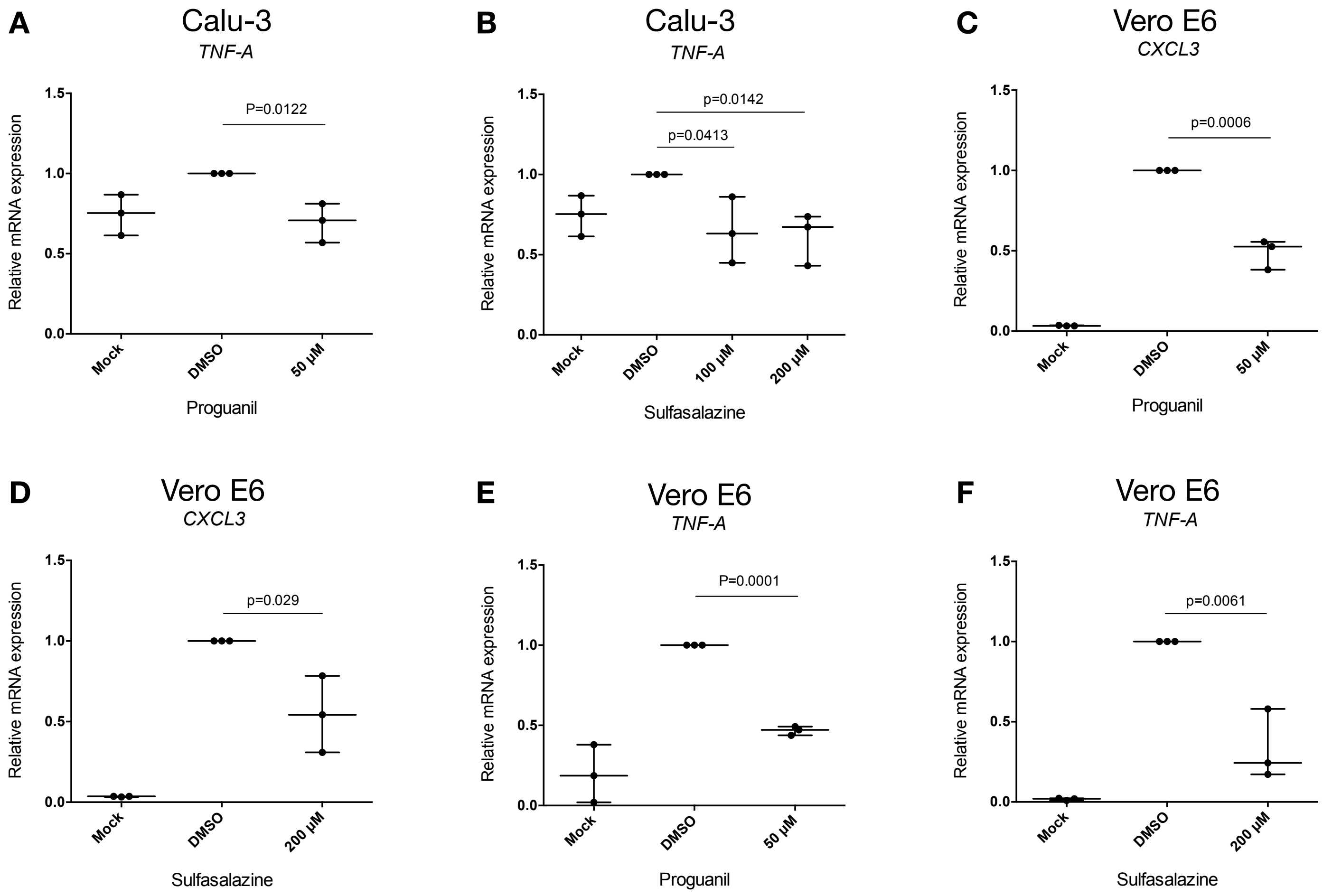


**Figure S8** Proguanil and Sulfasalazine significantly impair SARS-CoV-2 replication and blocked the p38/MAPK Signalling Activity. (A and B) RT-qPCR analysis of the indicated mRNAs from Calu-3 cells pre-treated with Proguanil or Sulfasalazine at indicated concentrations for three hours prior to infection with SARS-CoV-2 for 24 hours. Statistical test: Student's t test. Mean + S.D. of three independent replicates is shown. (C-F) RT-qPCR analysis of the indicated mRNAs from Vero E6 cells pre-treated with Proguanil or Sulfasalazine at indicated concentrations for three hours prior to infection with SARS-CoV-2 for 24 hours. Statistical test: Student's t test. Mean + S.D. of three independent replicates is shown.

Figure S9

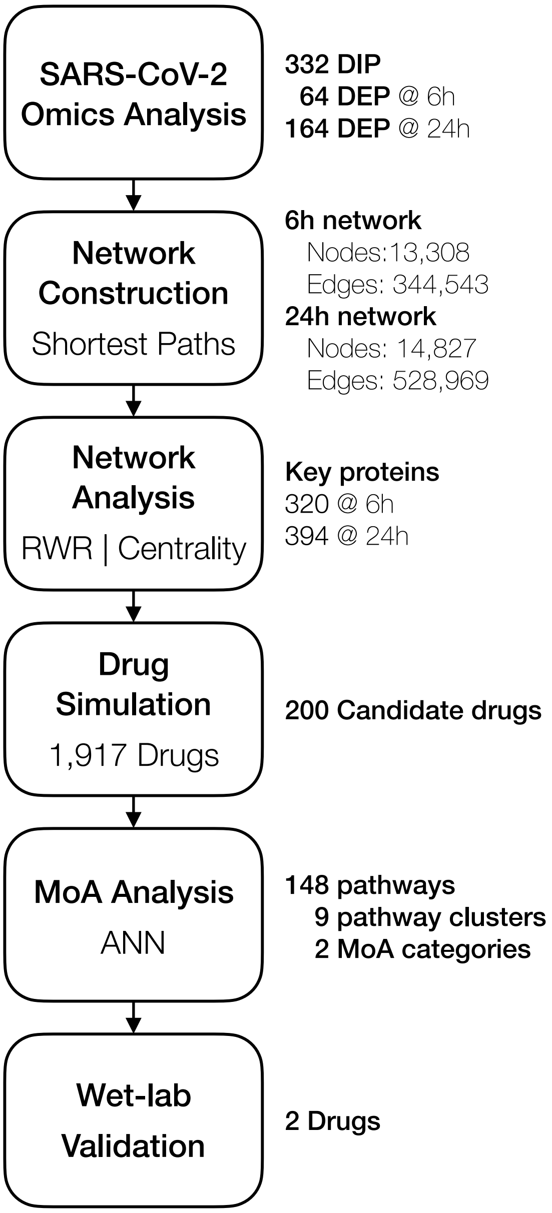


**Figure S9** Workflow

Table S1

**Table S1** Disease enrichment test result for key proteins in 6-hour network[1–3], 24-hour network[1–3]

### 6-hour

| Index | Name | P-value | Adjusted p-value |
|---|---|---|---|
| 1 | Arthritis | 1.265E-33 | 2.291E-30 |
| 2 | Lung disease | 2.703E-26 | 2.448E-23 |
| 3 | Toxic encephalopathy | 6.067E-23 | 3.662E-20 |
| 4 | Cerebrovascular disease | 2.757E-22 | 1.248E-19 |
| 5 | Brain disease | 6.893E-22 | 2.497E-19 |
| 6 | Hyperglycemia | 6.399E-21 | 1.932E-18 |
| 7 | Hypertension | 6.627E-18 | 1.715E-15 |
| 8 | Exanthem | 2.034E-13 | 4.604E-11 |
| 9 | Diabetic retinopathy | 4.275E-13 | 8.602E-11 |
| 10 | Pancreatitis | 4.551E-12 | 8.243E-10 |

### 24-hour

| Index | Name | P-value | Adjusted p-value |
|---|---|---|---|
| 1 | Brain disease | 1.308E-18 | 2.369E-15 |
| 2 | Arthritis | 5.054E-16 | 4.576E-13 |
| 3 | Toxic encephalopathy | 5.285E-15 | 3.190E-12 |
| 4 | Hyperglycemia | 3.395E-13 | 1.537E-10 |
| 5 | Cerebrovascular disease | 4.417E-13 | 1.600E-10 |
| 6 | Lung disease | 2.038E-11 | 6.151E-09 |
| 7 | Upper respiratory tract disease | 2.495E-11 | 6.456E-09 |
| 8 | Exanthem | 5.829E-11 | 1.319E-08 |
| 9 | Stomach cancer | 3.327E-09 | 6.694E-07 |
| 10 | Hypertension | 3.436E-09 | 6.223E-07 |

**Reference**


1. Gupta, A. *et al.* Extrapulmonary manifestations of COVID-19. *Nat. Med.* **26**, (2020).
2. Guan, W. *et al.* Clinical Characteristics of Coronavirus Disease 2019 in China. *N. Engl. J. Med.* **382**, 1708–1720 (2020).
3. Schett, G., Manger, B., Simon, D. & Caporali, R. COVID-19 revisiting inflammatory pathways of arthritis. *Nat. Rev. Rheumatol.* **16**, 465–470 (2020).

**Table S3** Examples of compounds in COVID-19 clinical trials

| Drug | MoA |
|---|---|
| **Captopril** | ACE2 inhibitor by Teva |
| **Chloroquine** | Anti-malarial drug by Sanofi |
| **Chloroquine**<br>**Deferoxamine**<br>**Methylprednisolone**<br>**Siltuximab** | Combination with Tocilizumab (immunosuppressive drug, mainly for the treatment of rheumatoid arthritis) preventing pneumonia by Genentech |
| **Dexamethasone** | Anti-inflammatory drug by Novartis |
| **Etoposide** | Cancer drug and also curing hemophagocytic lymphohistiocytosis that is classified as one of the cytokine storm syndromes due to virus infection by Pfizer |
| **Losartan** | ACE2 inhibitor by MSD |
| **Methylprednisolone**<br>**Sirolimus**<br>**Tacrolimus** | Immune-suppressant drugs by Pfizer |
| **Trimethoprim** | Combination antibiotics with Sulfamethoxazole preventing pneumocystis pneumonia and toxoplasmosis in people with HIV/AIDS by multiple pharma companies including Pfizer and GlaxoSmithKline |
